# Linked CD4^+^/CD8^+^ T cell neoantigen vaccination overcomes immune checkpoint blockade resistance and enables tumor regression

**DOI:** 10.1101/2023.05.06.539290

**Authors:** Joseph S. Dolina, Joey Lee, Spencer E. Brightman, Sara McArdle, Samantha M. Hall, Rukman R. Thota, Manasa Lanka, Ashmitaa Logandha Ramamoorthy Premlal, Jason A. Greenbaum, Ezra E.W. Cohen, Bjoern Peters, Stephen P. Schoenberger

## Abstract

Therapeutic benefit to immune checkpoint blockade (ICB) is currently limited to the subset of cancers thought to possess a sufficient tumor mutational burden (TMB) to allow for the spontaneous recognition of neoantigens (NeoAg) by autologous T cells. We explored whether the response of an aggressive low TMB squamous cell tumor to ICB could be improved through combination immunotherapy using functionally defined NeoAg as targets for endogenous CD4^+^ and CD8^+^ T cells. We found that, whereas vaccination with CD4^+^ or CD8^+^ NeoAg alone did not offer prophylactic or therapeutic immunity, vaccines containing NeoAg recognized by both subsets overcame ICB resistance and led to the eradication of large established tumors that contained a subset of PD-L1^+^ tumor-initiating cancer stem cells (tCSC), provided the relevant epitopes were physically linked. Therapeutic CD4^+^/CD8^+^ T cell NeoAg vaccination produced a modified tumor microenvironment (TME) with increased numbers of NeoAg-specific CD8^+^ T cells existing in progenitor and intermediate exhausted states enabled by combination ICB-mediated intermolecular epitope spreading. The concepts explored herein should be exploited for the development of more potent personalized cancer vaccines that can expand the range of tumors treatable with ICB.

## Introduction

Antibody-mediated blockade of PD-1 and CTLA-4 has been shown to therapeutically enhance pre-existing NeoAg-specific CD8^+^ T cell responses in several human tumors (1, 2). Recent findings support that clinical responsiveness to ICB strongly relies on a combination of overall TMB and a pre-treatment T-helper 1/interferon-γ (T_h_1/IFN-γ) inflammatory signature within a tumor (3–8). Of the majority of patients that do not respond to ICB, some are completely refractive (primary resistance) while others display a short-lived objective response followed by disease progression (secondary resistance) (9). Understanding and overcoming the relevant ICB resistance mechanisms in the non-responsive patient cohort would meaningfully increase both the therapeutic index and number of patients who could benefit from this important treatment paradigm.

Although endogenous major histocompatibility complex class I (MHC I)-restricted T cell responses appear critical for the benefit of ICB, the durability of CD8^+^ T cells once inside the immunosuppressive TME is questionable (10). In both clinical and preclinical settings, emerging evidence has revealed that neoplastic cells can evade ICB-mediated immune responses indirectly through immunoediting, a process by which the immunogenicity of tumor cells is reduced via downregulation of presented NeoAg (5, 9). tCSC, which are critical for tumor formation and growth, are particularly difficult to eradicate and display intrinsic resistance to chemotherapy and ICB in part due to their slower growth rate and elevated expression of the ligands for PD-1 and CTLA-4 inhibitory receptors (PD-L1 and CD80, respectively) (11–13). Further, tumor cells can directly promote the formation of ICB-refractory exhausted CD8^+^ T cells (T_ex_) prior to treatment and exhaustion of effector cells after immunotherapeutic reinvigoration via persistent NeoAg display (similar to chronic viral infection) or secretion of immunosuppressive factors such as vascular endothelial growth factor-A (VEGF-A) (14–19). In addition, low levels of NeoAg expression and reduced MHC I affinity can also result in poor CD8^+^ T cell priming and actively drive exhaustion (8, 14, 20). Each of these mechanisms can limit the efficacy of endogenous CD8^+^ T cell responses mobilized by ICB.

NeoAg have emerged as the targets of successful immunotherapy in a number of clinical settings including adoptive cellular transfer, ICB, and personalized vaccines (21). As such, there is significant interest in identifying and exploiting the subset of expressed mutations by which a tumor can be recognized by autologous T cell responses. Most of these involve analysis of peptides containing mutations for predicted binding to MHC I molecules, thereby confining the vaccine-induced T cell responses to the CD8^+^ T cell subset, despite the fact that MHC class II (MHC II)-restricted CD4^+^ T cells have been only recently demonstrated to potentiate anti-tumor immunity through a variety of mechanisms including providing T cell help to CD8^+^ T cells via CD40-mediated activation of antigen presenting cells (APC), locally producing IL-21 to directly sustain CD8^+^ T cell effector activity, and as direct CD4^+^ T effectors (22–30). Although CD4^+^ T cells are found to be necessary for sustaining high avidity tumor-specific CD8^+^ T cell responses (31–36), it is unclear if and how this extends to established ICB-resistant neoplastic disease where low to moderate avidity CD8^+^ T_ex-term_ typically dominate the TME and whether natural CD4^+^ T cell tumor specificity is needed for the immunotherapeutic treatment of MHC II^−^ tumors in this context (20, 23, 25, 37–39).

Rather than relying on prediction, we sought to utilize a functional approach to NeoAg identification based on monitoring physiological CD4^+^ and CD8^+^ T cell responses to tumor-derived antigens. These results show that natural NeoAg recognized by both CD4^+^ and CD8^+^ T cells are superior compared to epitopes priming either cell type alone. Remarkably, CD4^+^ T cell target antigen did not need to be tumor-restricted in both prophylactic and therapeutic settings and could instead be targeted to a universal MHC II-binding helper epitope. Lastly, we demonstrate that ICB-resistance can effectively be overcome by combination with NeoAg vaccination in a synergistic mechanism to sustain stable CD8^+^ T cell responses capable of resisting the onset of terminal exhaustion and targeting both PD-L1^+^ and PD-L1^−^ tumor cells.

## Results

### Cancer cell stemness and intrinsic resistance mechanisms

Squamous cell carcinoma VII (SCC VII) is a spontaneously arising MHC II^−^ murine tumor which closely resembles human head and neck squamous cell carcinoma (HNSCC) in several key features including pulmonary and lymph node (LN) metastasis, poor immunogenicity, and, importantly, resistance to chemotherapeutic and immunotherapeutic intervention (40–44). Several lines of evidence suggest that a small fraction of tCSC marked by elevated CD44 expression exist within this class of neoplasm linked with baseline tumor growth, metastasis, and resistance mechanisms (11, 45, 46). In this study, we initially noted that the SCC VII transcriptome shared several common signaling pathways with various human cancer types (including HNSCC) in aligned NCBI OncoGEO tumor datasets featuring significant similarity in Rho kinase signaling, which is critical in governing tCSC formation and maintenance (Supplemental Figure 1A) (47, 48). Both murine SCC VII and primary human tumor cells collected from HNSCC patients responded similarly in vitro to Rho kinase inhibition, which promoted a CD44^hi^ tCSC phenotype additionally associated with co-expression of other stem cell markers including ALDH1A1, EpCAM, and EGFR (Supplemental Figure 1, B-D) (11). SCC VII tCSC also displayed cardinal features of invasive human tCSC including impairments in actin stress fiber formation (Supplemental Figure 2A) and more rapid migration in a wound closure assay (Supplemental Figure 2B). In the absence of disrupting Rho kinase activity, we observed that early passage SCC VII contained a CD44^hi^ subpopulation that became absent over time as cells were passaged in basal media in vitro (Supplemental Figure 3A). In addition, late passage SCC VII lacking CD44^hi^ tCSC failed to form tumors in vivo (Supplemental Figure 3B). Thus, SCC VII appears to closely mirror the cellular heterogeneity commonly observed in human HNSCC.

We next characterized common non-immune and immune resistance mechanisms deployed by CD44^lo^ versus CD44^hi^ SCC VII in vivo. To establish the inherent chemoresistance of these subsets, SCC VII-Luc/GFP tumors were grown in groups of mice for 10 days, and cell death was assessed by measuring active caspase-3 in tumor cells 7 days after saline versus a maximum tolerated dose of cisplatin delivered intraperitoneally (IP). CD44^hi^ tCSC had lower overall active caspase-3 compared to CD44^lo^ tumor cells in saline-treated mice. After one treatment with cisplatin, the amount of active caspase-3 significantly increased in CD44^lo^ tumor cells while CD44^hi^ tCSC remained unresponsive (Supplemental Figure 3, C and D). These results suggest that CD44^hi^ tCSC have an inherent increased chemoresistance compared to more differentiated cells. Additionally, consistent with the immunosuppressive phenotype of human HNSCC-derived CD44^hi^ tCSC, CD44^hi^ SCC VII tCSC had significantly elevated PD-L1 expression compared to CD44^lo^ cells in the absence or presence of strong T_h_1 inflammation following delivery of 50 μg polyinosinic-polycytidylic acid (polyI:C), a synthetic Toll-like receptor 3 (TLR3) ligand (Supplemental Figure 3E) (12). In total, these results reveal that SCC VII tumors may be difficult to treat in situ by conventional standard of care therapies given to HNSCC patients (chemotherapy and/or ICB) due to the inherent resistance mechanisms deployed by stem versus differentiated cells.

### Functional identification of neoantigens based on endogenous CD4^+^ and CD8^+^ T cell reactivity

In pursuit of NeoAg targets of natural immune responses against SCC VII, we first established its inherent immunogenicity. C3H/HeJ mice were subcutaneously (SC) immunized with 1×10^7^ irradiated SCC VII cells, either alone or supplemented with 50 μg polyI:C. Immunized mice were challenged 14 days later with 5×10^5^ live SCC VII cells transduced to express luciferase and green fluorescent protein (SCC VII-Luc/GFP) to enable tracking by bioluminescence (BLI). Whereas whole-cell vaccination with irradiated SCC VII alone did not protect mice from tumor outgrowth following challenge, revealing SCC VII as a poorly-immunogenic tumor by this classical definition, prophylaxis was achievable through co-delivery of polyI:C (Figure 1, A and B). Thus, SCC VII contains antigens capable of conferring protective immunity. This depends on both CD4^+^ and CD8^+^ T cells, as depletion of either subset before or after vaccination led to tumor outgrowth following subsequent challenge (Figure 1, C and D). Notably, tumors in mice depleted of CD4^+^ T cells just prior to challenge displayed a reduced growth rate compared to controls, suggesting that this subset is required at both the initiation phase of the vaccine-induced response and later to maintain its efficacy following challenge.

**Figure 1.**
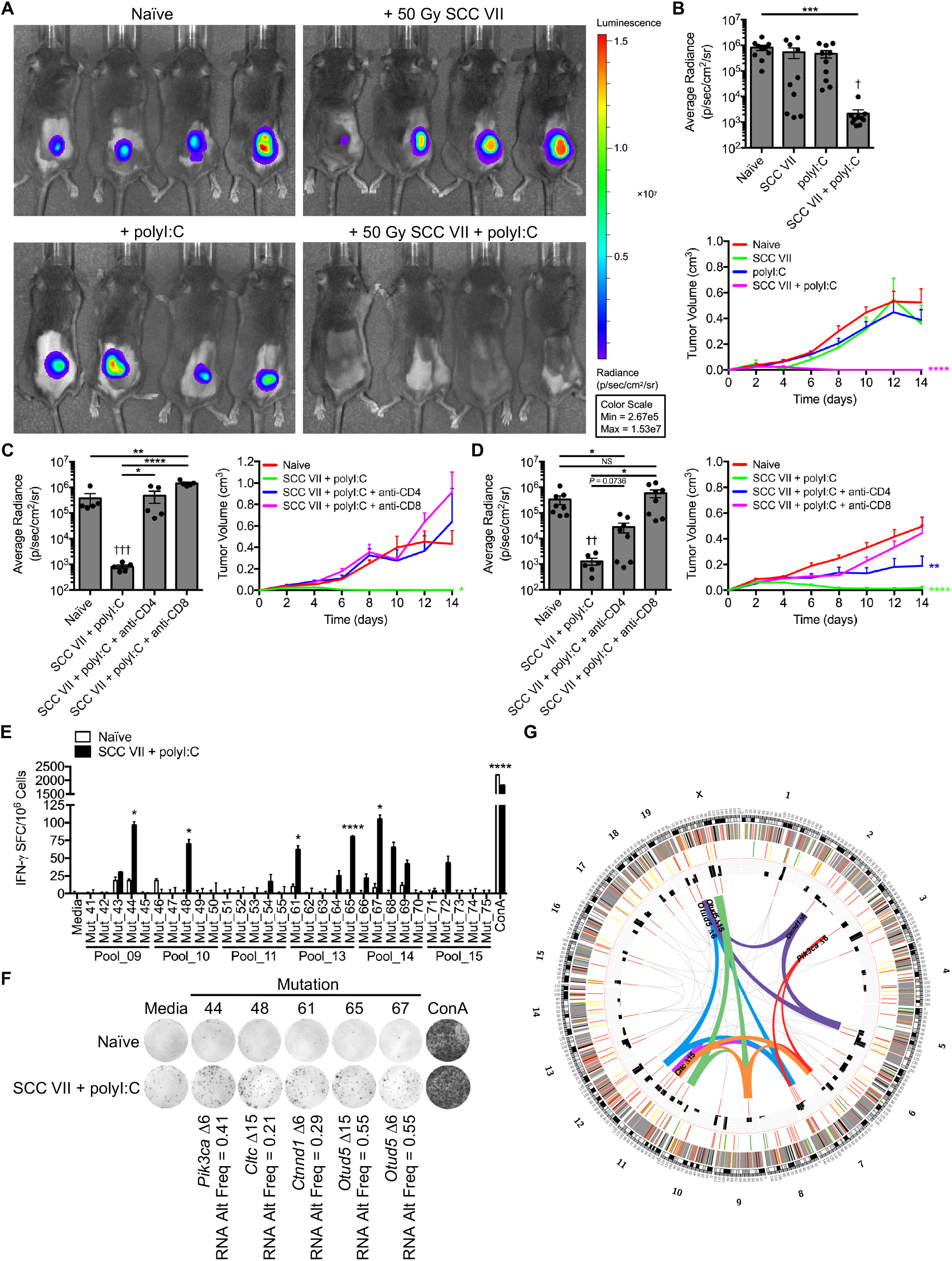
SCC VII and polyI:C co-immunization elicits protection from live tumor challenge and subsequent neoantigen identification. (**A**-**D**) C3H/HeJ mice immunized with 1×10^7^ irradiated SCC VII cells, 50 μg polyI:C, or both in combination and subsequently challenged with 5×10^5^ live SCC VII-Luc/GFP cells 14 days later. (**A** and **B**) Bioluminescence of mice bearing SCC VII-Luc/GFP tumors 14 days after challenge and recorded tumor volume kinetics (n = 10 per group). Mice depleted of CD4^+^ or CD8^+^ cells (**C**) prior to or (**D**) after co-immunization with irradiated SCC VII cells and polyI:C assessed for day 14 SCC VII-Luc/GFP bioluminescence and tumor volume kinetics (n = 5-8 per group). (**E** and **F**) Groups of naïve and immunized/challenged C3H/HeJ mice as in (**A**-**D**) assessed for the presence of IFN-γ-producing splenic and Ig LN mononuclear cells at day 28 via ELISPOT after re-stimulation with NeoAg-pulsed BMDCs (n = 3 per group). (**G**) Circos plot representative of total and filtered mutations identified from Exome-Seq and RNA-Seq of SCC VII, selected peptides, and pooled/single peptide IFN-γ ELISPOT results. Outside to inside tracks are arranged as 1.) chromosome with Mb labels of physical distance, 2.) somatic mutations, 3.) somatic strict mutations, 4.) VAF, and 5.) selected peptides. Inner region summarizes significant IFN-γ ELISPOT results from (**E** and **F** and Supplemental Figure 4) where ribbons and gene names represent peptide pools or individual peptides, respectively. Pool_9 (red), Pool_10 (orange), Pool_11 (green), Pool_13 (purple), Pool_14 (blue), and Pool_15 (magenta). Size of the ribbons or gene names correlates with the number of SFC. All experiments were performed two or more times and data indicate means ± s.e.m.; (**B**-**D**, bioluminescence) \**P* < 0.05, \*\**P* < 0.01, \*\*\**P* < 0.001, and \*\*\*\**P* < 0.0001 (Student’s t test); ^†^*P* < 0.05, ^††^*P* < 0.01, and ^†††^*P* < 0.001 (one-way ANOVA and Dunnett’s post hoc test relative to naïve); (**B**-**D**, tumor volume) \**P* < 0.05, \*\**P* < 0.01, and \*\*\*\**P* < 0.0001 (two-way ANOVA and Dunnett’s post hoc test relative to naïve); (**E**) \**P* < 0.05 and \*\*\*\**P* < 0.0001 (Student’s t test of data with SI > 2 and Poisson < 5%).

To identify SCC VII antigens conferring protective immunity, we employed an approach which combined genomic sequencing to detect well-expressed coding mutations with functional analysis of natural immune responses to tumor antigens. The SCC VII tumor exome was compared to that of normal control C3H/HeJ caudal tissue samples. This analysis yielded 1,481 variants in coding sequences among 4,771 total variants detected in the tumor versus reference exome. Of these, 270 could be confirmed as expressed by at least one read of the variant base in the tumor RNA, with 39 mutations reaching our selected expression threshold of 20% variant allele frequency (VAF) and ≥ 10 reads in the tumor RNA sample. These 39 mutations were translated into amino acid sequences, and 20-mer peptide pairs were synthesized for each mutation in which the mutated amino acid was placed at position 6 or 15 (or position 10 in one case involving insufficient amino acids near an alternative splicing site) within the linear peptide flanked by wild type sequence (Supplemental Table 1).

The 81 candidate peptides representing the 39 filtered mutations were tested as targets for T cells generated by immunization with the irradiated SCC VII ± polyI:C and live tumor challenge protocol described above. This involved re-stimulation of splenic and tumor-draining inguinal LN (Ig LN) mononuclear cells isolated 14 days after challenge with bone marrow-derived dendritic cells (BMDC) pulsed with 16 pools of 20-mer peptides in ELISPOT assays for assessment of IFN-γ effector cytokine production. Significant frequencies of IFN-γ spot forming cells (SFC) over background were found for 6 of the 16 peptide pools screened (Supplemental Figure 4, A and B). Peptide pools that produced strong IFN-γ responses were subsequently deconvoluted to detect the specific mutant peptides targeted. This analysis revealed *Pik3ca* (Mut_44), *Cltc* (Mut_48), *Ctnnd1* (Mut_61), and *Otud5* (Mut_65 and Mut_67) as the mutated genes recognized by natural immune responses to SCC VII (Figure 1, E and F). Positive responses observed for Mut_65 and Mut_67, which contain the same mutation in the *Otud5* deubiquitinase gene (at positions 15 or 6 within the 20-mer peptide, respectively), served as an internal control for our in vitro assay when compared to the absence of responses against Mut_64 and Mut_66, which contain the same nucleotide change but result in a different peptide product due to nearby alternative splicing. Mut_44 (*Pik3ca* Δ6) corresponds to a T1025A modification in the catalytic domain of phosphatidylinositol 3-kinase—recently identified as a novel driver mutation in addition to the dominant H1047R affecting the same domain in human cancers (49). Mut_48 (*Cltc* Δ15) maps to the propeller domain of the clathrin heavy chain known to support both mitosis and nutrient uptake by cancer cells (50, 51). Mut_61 (*Ctnnd1* Δ6) is located nearby the ARM domain of catenin δ-1 as I489N—also documented as a driver mutation affecting cell adhesion (Supplemental Figure 5, A-D) (52). The steps involved in identification of somatic variants, selection of candidate mutations for functional testing, and functional validation of NeoAg are graphically represented as a Circos plot (Figure 1G and Supplemental Table 2).

The immunogenicity of the four SCC VII NeoAg was next investigated. C3H/HeJ mice were immunized SC once or boosted three weeks later with a pool of the five recognized 20-mer peptides + polyI:C. Mice were challenged 10 days after the last (booster) vaccination with live SCC VII-Luc/GFP SC on the opposite flank, and tumor outgrowth was subsequently monitored by BLI and caliper measurements. Whereas a single injection of the pooled NeoAg peptides did not protect from SCC VII tumor challenge, boosting this response with a second immunization led to significantly smaller tumor sizes at all time points assayed (Supplemental Figure 6, A-D). CD4^+^ and CD8^+^ T cells were critical for mediating the protective immunity elicited by the NeoAg vaccine, as this was lost with depletion of either population prior to challenge (Supplemental Figure 7, A and B). When the individual NeoAg peptides were tested for their contribution to the observed immunity, only Mut_48 (*Cltc* Δ15) demonstrated the ability to confer protection from challenge (Figure 2A), while the wild type peptide (WT_48) was not protective (Figure 2B). These data indicate that the T cell response to Mut_48 mediates protective immunity following prophylactic peptide vaccination.

**Figure 2.**
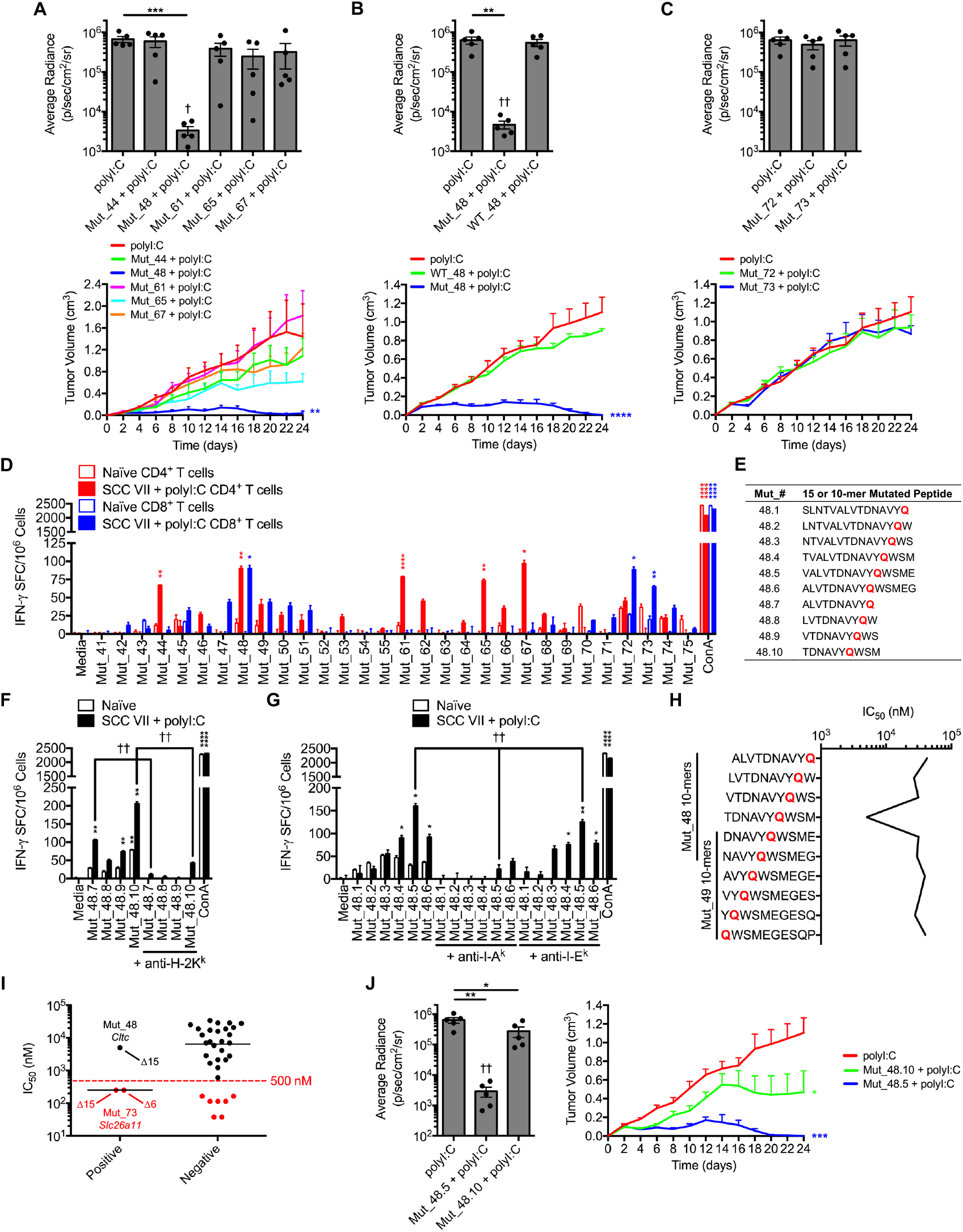
Deconvolution of CD4^+^ and CD8^+^ T cell responses to SCC VII-derived neoantigens. (**A**-**C**) C3H/HeJ mice vaccinated with 50 μg polyI:C alone or in combination with 5 μg solubilized 20-mers in a booster regimen 21 days apart. All sets of mice were challenged with 5×10^5^ live SCC VII-Luc/GFP cells 31 days after primary vaccination. Individual (**A**) Mut_44, Mut_48, Mut_61, Mut_65, or Mut_67 long peptides, (**B**) Mut_48 versus WT_48, and (**C**) Mut_72 and Mut_73 long peptides reported as bioluminescence of mice at 14 days after challenge and tumor volume kinetics (n = 6 per group). (**D**) Groups of naïve C3H/HeJ mice compared to animals that received a 1×10^7^ irradiated SCC VII cell and 50 μg polyI:C immunization and later challenge assessed for the presence of IFN-γ-producing CD4^+^ and CD8^+^ T cells sorted from spleens and Ig LNs at day 28 post-immunization via ELISPOT after re-stimulation with NeoAg-pulsed BMDCs (n = 3 per group). (**E**) 15- and 10-mer minimal peptides unique to the parent Mut_48 20-mer peptide with shifted H129Q mutation positions. (**F** and **G**) Splenic and Ig LN (**F**) CD8^+^ T cells and (**G**) CD4^+^ T cells sorted from vaccinated mice co-cultured with Mut_48-derived minimal peptide-pulsed BMDCs for quantification of IFN-γ-producing cells via ELISPOT. Blocking antibodies against I-A^k^, I-E^k^, and H-2K^k^ included in select wells (n = 3 per group). (**H**) MHC I predictions of minimal peptide binding to murine H-2K^k^ using the IEDB NetMHCpan (v4.0) tool. (**I**) CD8^+^ T cell ELISPOT responses against all peptides in Pool_9, Pool_10, Pool_11, Pool_13, Pool_14, and Pool_15 clustered as eliciting robust IFN-γ production (positive) versus those that did not (negative). Represented are MHC I predictions of minimal peptide binding to murine H-2K^k^ using the IEDB NetMHCpan (v4.0) tool where > 500 nM (black) and < 500 nM (red) is noted. Designated on the positive response partition is the corresponding mutant peptide. (**J**) Day 14 bioluminescence and tumor volume kinetics of C3H/HeJ mice challenged with SCC VII-Luc/GFP tumors after a booster immunization as in (**A**) with Mut_48.5 and Mut_48.10 minimal peptides (n = 6 per group). All experiments were performed two or more times and data indicate means ± (**A**-**G** and **J**) s.e.m. or (**I**) median; (**A**-**C** and **J**, bioluminescence) \**P* < 0.05, \*\**P* < 0.01, and \*\*\**P* < 0.001 (Student’s t test); ^†^*P* < 0.05 and ^††^*P* < 0.01 (one-way ANOVA and Dunnett’s post hoc test relative to polyI:C); (**A**-**C** and **J**, tumor volume) \**P* < 0.05, \*\**P* < 0.01, \*\*\**P* < 0.001, and \*\*\*\**P* < 0.0001 (two-way ANOVA and Dunnett’s post hoc test relative to polyI:C); (**D**, **F**, and **G**) \**P* < 0.05, \*\**P* < 0.01, and \*\*\*\**P* < 0.0001 (Student’s t test of data with SI > 2 and Poisson < 5%); (**F**) ^††^*P* < 0.01 (Student’s t test); (**G**) ^††^*P* < 0.01 (one-way ANOVA and multiple comparison Tukey’s post hoc test).

### Effective vaccination requires both MHC I- and II-presented neoantigens

We next determined the T cell subsets involved in the natural NeoAg-specific immune response by assessing the reactivity of CD4^+^ versus CD8^+^ T cells isolated from mice immunized with SCC VII tumor cells. We found that Mut_48 was recognized by both CD4^+^ and CD8^+^ T cells, whereas Mut_44, Mut_61, Mut_65 and Mut_67 were solely recognized by CD4^+^ T cells. In addition, we found that isolated CD8^+^ T cells recognized Mut_72 and Mut_73, distinct peptides containing the same missense mutation in the *Slc26a11* gene (Figure 2D). However, neither Mut_72 and Mut_73 were capable of conferring protective immunity against SCC VII in vivo following prime/boost vaccination (Figure 2C). These results collectively demonstrate that only Mut_48, which is recognized by both CD4^+^ and CD8^+^ T cells, is capable of inducing effective prophylactic immunity through peptide vaccination.

To determine whether the epitopes recognized by each subset were identical or distinct, we used IFN-γ ELISPOT to quantitate the relative response magnitude to a panel of 10- and 15- mer peptides, designated Mut_48.1-Mut_48.10, containing the Mut_48 H129Q mutation (Figure 2E). CD8^+^ T cells isolated from SCC VII/polyI:C-immunized mice produced the greatest amount of IFN-γ upon recognition of the Mut_48.10 10-mer with antibody blockade demonstrating its presentation by H-2K^k^ (Figure 2F). Co-purified CD4^+^ T cells showed the greatest reactivity to the Mut_48.5 15-mer presented via I-A^k^ (Figure 2G) and did not react with any designed 10-mer as expected (data not shown). Further, in silico prediction of Mut_48-derived 10-mer binding to H-2K^k^ using the NetMHCpan (v4.0) (53) method estimated poor affinities for most peptides, with the best affinity predicted for Mut_48.10 at 4988.7 nM IC_50_. Notably, the 10-mer peptides contained within the Mut_48.7-48.10 series elicited IFN-γ production despite predicted H-2K^k^ binding IC_50_ values being above the 500 nM cutoff used for most screening protocols (Figure 2H). These results further confirmed that Mut_48 contains a CD8^+^ T cell minimal epitope, Mut_48.10, within the longer CD4^+^ T cell epitope, Mut_48.5, thereby endowing IFN-γ production from both T cell subsets. An expanded analysis of H-2K^k^ binding predictions for IFN-γ stimulatory Mut_72 and Mut_73 resulted in 250.9 nM IC_50_ affinities for both peptides suggesting that the Immune Epitope Database and Analysis Resource (IEDB) NetMHCpan (v4.0) tool algorithm is ∼66% efficient at filtering for in vitro immunoreactivity of CD8^+^ T cells (Figure 2I). The unbiased functional approach utilized in this study to identify NeoAg therefore allowed us to more efficiently probe both CD4^+^ and CD8^+^ T cell epitopes in the same assay system.

Immunization studies showed that the Mut_48.5 15-mer, containing both CD4^+^ and CD8^+^ T cell-recognized minimal epitopes, was protective against live SCC VII cell challenge in vivo to a degree comparable to the Mut_48 20-mer. Additionally, the CD4^+^ T cell epitope was entirely necessary for the observed protection as immunization of mice with the truncated Mut_48.10 10-mer containing only the CD8^+^ T cell epitope was partially protective (Figure 2J). These findings are reminiscent of prior animal studies and clinical trials where vaccination with CD8^+^ T cell NeoAg alone results in a detectable response and tumor regression, but eventual tolerance and later relapse (28).

### Tumor specificity of CD4^+^ T cells and provision of help to CD8^+^ T cells

We sought to determine if the help provided by the CD4^+^ T cell response was strictly tumor-specific. Mice were immunized with the Mut_48.10 CD8^+^ T cell minimal epitope mixed with either the SCC VII-derived Mut_44 (*Pik3ca* Δ6), which is recognized by CD4^+^ T cells (Figure 2D) or the pan-DR epitope peptide (PADRE[X] where X = cyclohexylalanine in the third position). PADRE(X) is an immunogenic peptide originally designed for broad specificity to human DR MHC II molecules but is also capable of providing T cell help to antigen-specific CD8^+^ cytotoxic T lymphocyte (CTL) responses in C57BL/6 mice in vivo and can competitively bind to C3H/HeJ I-A^k^ with high affinity in vitro (54). However, CD4^+^ T cells activated by PADRE(X) would be unable to contribute to anti-tumor immunity by any mechanism requiring tumor specificity after relaying T cell help. Regardless of the origin of the helper epitope, co-delivery of functional CD4^+^ T cell antigens improved Mut_48.10-mediated prophylactic immunity to a similar degree as the full Mut_48 20-mer containing both CD4^+^ and CD8^+^ T cell epitopes. Further, covalent linkage of CD4^+^ and CD8^+^ T cell antigens via a triple alanine repeat (-AAA-) resulted in superior protection against SCC VII challenge (Figure 3). The efficacy of Mut_48 is thus related to presentation of the CD4^+^ T cell helper antigen alongside a tumor-specific CTL NeoAg most likely by the same APC, mechanistically consistent with T_h_-mediated ‘licensing’ of APC to program optimal CD8^+^ T cell responses (23, 26). Furthermore, these results suggest that other effector functions of CD4^+^ T cells requiring tumor specificity are dispensable in this model.

**Figure 3.**
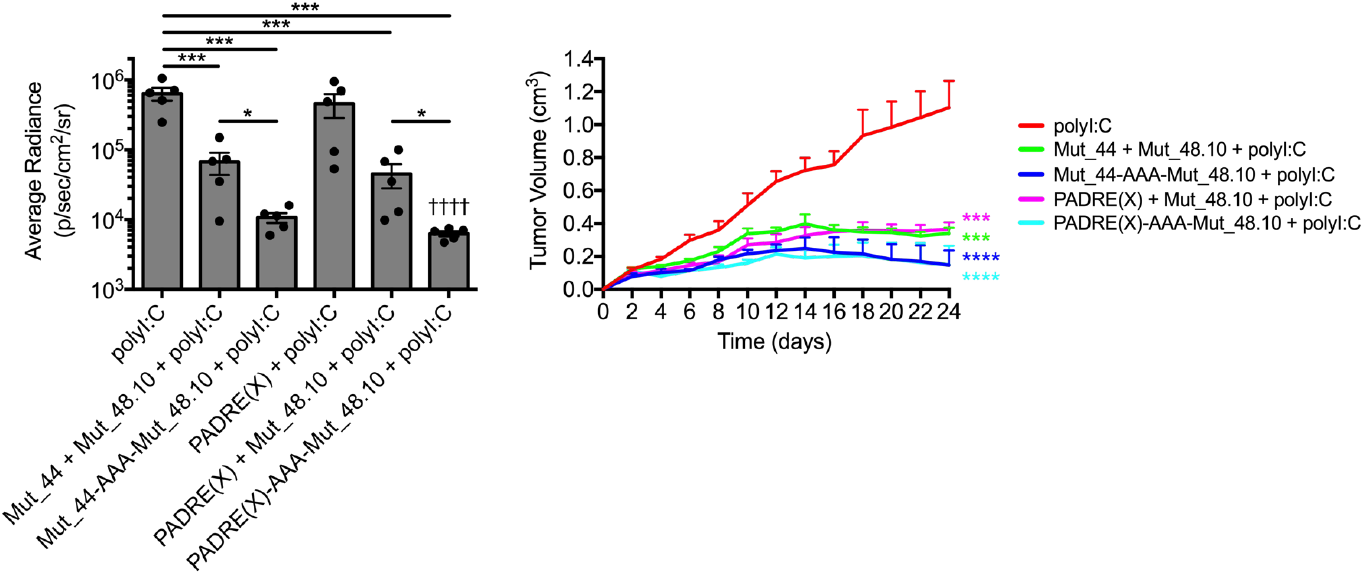
Tethered CD4^+^ T cell helper and minimal CD8^+^ T cell epitope vaccines lead to maximal SCC VII tumor growth inhibition. C3H/HeJ mice vaccinated with 50 μg polyI:C alone or in combination with 5 μg solubilized PADRE(X) or Mut_44 long-mers untethered or tethered to the Mut_48.10 minimal epitope in a booster regimen 21 days apart. All sets of mice were challenged with 5×10^5^ live SCC VII-Luc/GFP cells 31 days after primary vaccination. Day 14 bioluminescence and tumor volume kinetics of challenged C3H/HeJ mice. All experiments were performed two or more times and data indicate means ± s.e.m.; (bioluminescence) \**P* < 0.05, and \*\*\**P* < 0.001 (Student’s t test); ^††††^*P* < 0.0001 (one-way ANOVA and Dunnett’s post hoc test relative to polyI:C); (tumor volume) \*\*\**P* < 0.001 and \*\*\*\**P* < 0.0001 (two-way ANOVA and Dunnett’s post hoc test relative to polyI:C).

### Therapeutic resistance to immune checkpoint blockade is overcome by combination with neoantigen vaccination

ICB monotherapy has been shown to amplify endogenous NeoAg-specific T cell responses and generate de novo NeoAg responses when combined with prediction-based vaccines leading to an increase in progression-free survival in patients (1, 2, 55, 56). We therefore assessed whether a NeoAg vaccine based on validated targets could be rationally combined with this strategy. NeoAg-specific T cell responses were first induced through immunization with the pooled NeoAg vaccine (targeting the *Pik3ca*, *Cltc*, *Ctnnd1*, and *Otud5* mutations) using the prime/boost protocol described above. Three days after challenge with live SCC VII-Luc/GFP tumors, mice received blocking antibodies to either PD-1 or CTLA-4 by IP injection, and the effect on tumor outgrowth was measured. In both cases, ICB significantly accelerated the ability of the NeoAg vaccine to mediate therapeutic immunity against the growing SCC VII tumors compared to ICB alone. We found that combining ICB with peptide vaccine prevented the late phase (> day 24) relapse of SCC VII tumors observed in ∼50% of C3H/HeJ mice receiving NeoAg vaccination alone (Figure 4, A and B and Supplemental Figure 8). Elimination of palpable tumors was notably hastened with combinatorial anti-PD-1 and NeoAg vaccination with a synergistic effect apparent at day 14 during the early kinetic phase of active rejection (Figure 4A). Further, we examined memory T cell responses at day 42 post-challenge by IFN-γ ELISPOT against the NeoAg vaccine and found that PD-1 blockade increased the magnitude of Mut_48-specific T cell responses and showed evidence of intermolecular epitope spreading to Mut_72 and Mut_73 (Figure 4, C and D), targets that were not included in the peptide vaccination but had previously been observed to elicit CD8^+^ T cell responses upon physical separation from CD4^+^ T cells (Figure 2D). Anti-CTLA-4 treatment, in contrast, did not display synergy nor significantly affect the absolute number of Mut_48-specific T cells and exhibited reduced epitope spreading to other specificities (Figure 4B, Supplemental Figure 8, and Supplemental Figure 9, A and B).

**Figure 4.**
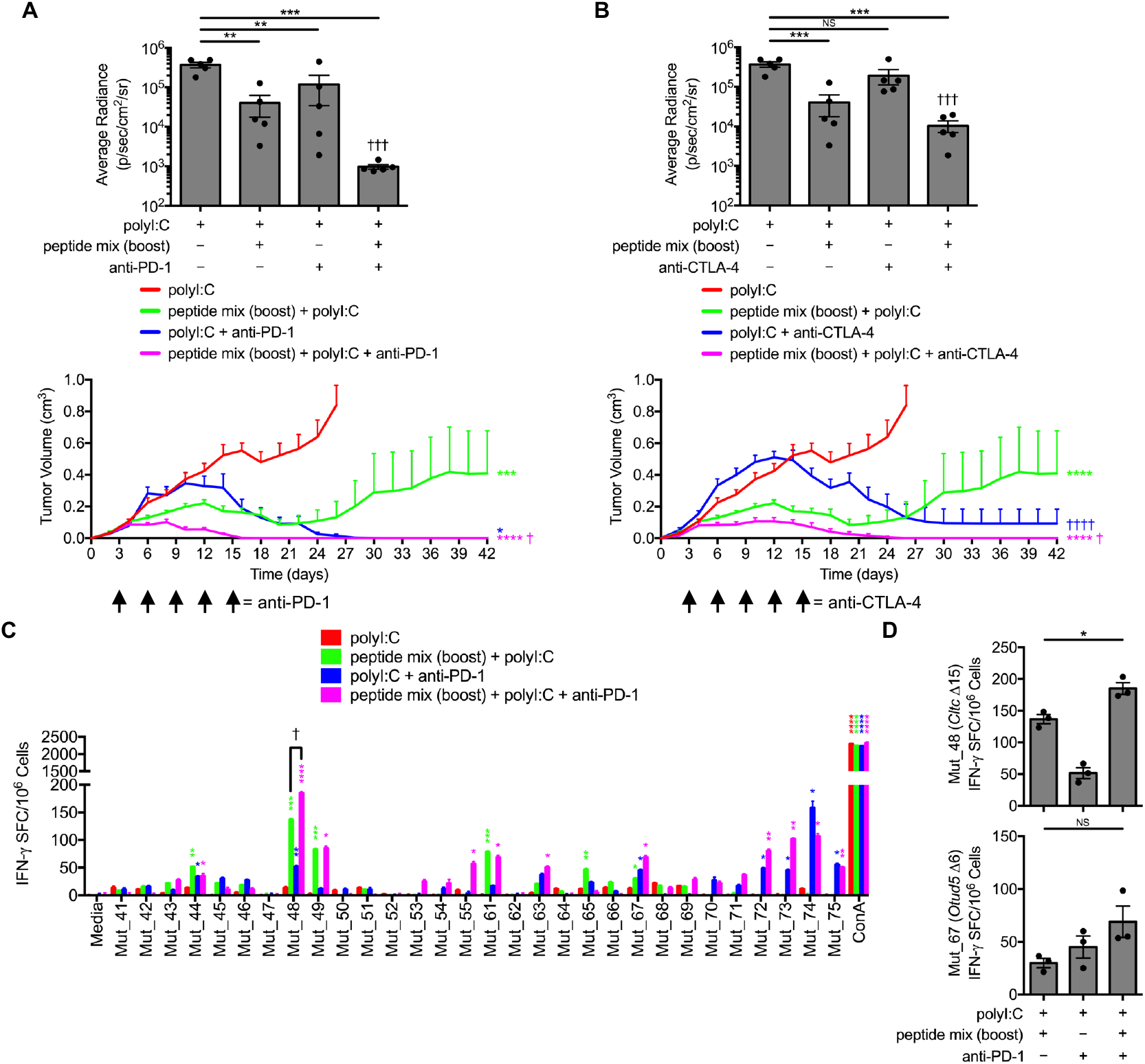
Anti-PD-1 checkpoint blockade additively increases *Cltc* Δ15-specific memory frequency and promotes dominant intermolecular epitope spreading. C3H/HeJ mice vaccinated with 50 μg polyI:C alone or in combination with prime/boost regimens of a 5×5 μg mixture containing solubilized Mut_44, Mut_48, Mut_61, Mut_65, and Mut_67 long peptides. All groups of mice were challenged with 5×10^5^ live SCC VII-Luc/GFP cells 31 days after primary vaccination. Treatment with (**A**) anti-PD-1 or (**B**) anti-CTLA-4 therapeutically began at day 3 after tumor cell inoculation (arrow) with bioluminescence of mice bearing live tumors at 14 days after challenge (upper panels) and tumor volume kinetics (lower panels) (n = 6 per group). (**C** and **D**) Splenic and Ig LN mononuclear cells isolated at day 42 from anti-PD-1-treated groups and controls assessed for IFN-γ-production via ELISPOT after re-stimulation with NeoAg-pulsed BMDCs (n = 3 per group). All experiments were performed two or more times and data indicate means ± s.e.m.; (**A** and **B**, bioluminescence) \*\**P* < 0.01 and \*\*\**P* < 0.001 (Student’s t test); ^†††^*P* < 0.001 (one-way ANOVA and Dunnett’s post hoc test relative to polyI:C); (**A** and **B**, tumor volume) \**P* < 0.05, \*\*\**P* < 0.001, and \*\*\*\**P* < 0.0001 (two-way ANOVA and Dunnett’s post hoc test relative to polyI:C); ^†^*P* < 0.05 and ^††††^*P* < 0.0001 (two-way ANOVA and Dunnett’s post hoc test relative to peptide mix boost + polyI:C); (**C**) \**P* < 0.05, \*\**P* < 0.01, \*\*\**P* < 0.001, and \*\*\*\**P* < 0.0001 (Student’s t test of data with SI > 2 and Poisson < 5%); ^†^*P* < 0.05 (Student’s t test); (**D**) \**P* < 0.05 (Student’s t test).

Given the synergistic potency of combining PD-1 blockade with NeoAg peptide vaccination, and the ability of the Mut_48 peptide to induce both CD4^+^ and CD8^+^ T cell responses against the SCC VII tumor, we examined whether these could be combined to treat large established tumors. SCC VII-Luc/GFP tumors were grown in groups of mice and allowed to reach a volume of ∼300-400 mm^3^ before treatment with two cycles of contralateral SC Mut_48 + polyI:C mixtures and/or IP anti-PD-1 on days 10 and 24. The Mut_48 vaccine alone did not result in a therapeutic benefit, whereas anti-PD-1 displayed varying degrees of primary and secondary resistance, only sometimes leading to initial tumor control that was subsequently lost. In contrast, combining PD-1 blockade with the Mut_48 NeoAg vaccine resulted in the complete and durable (> 90 days) eradication of large established tumors (Figure 5, A-D). In vitro re-stimulation of lymphocytes from the spleen and tumor-draining Ig LNs revealed combining NeoAg and anti-PD-1 treatments resulted in a synergistic boosting of memory phase Mut_48-specific T cell responses (Figure 5E). Further, NeoAg and anti-PD-1 co-administration significantly increased the number of total CD8^+^ T cells within tumor-infiltrating lymphocyte (TIL) fractions when isolated at the day 17 effector phase after the first round of immunotherapy, whereas conventional (T_conv_) and regulatory (T_reg_) CD4^+^ T cell numbers remain unchanged (Figure 5F). We additionally noted that rejection of tumor by Mut_48 and anti-PD-1 treatments was also accompanied by inhibition of SCC VII metastasis to regional LN (Figure 5G). These data suggest that functional NeoAg-mediated tumor rejection and prevention of regional metastasis is therapeutically optimal after combination with ICB.

**Figure 5.**
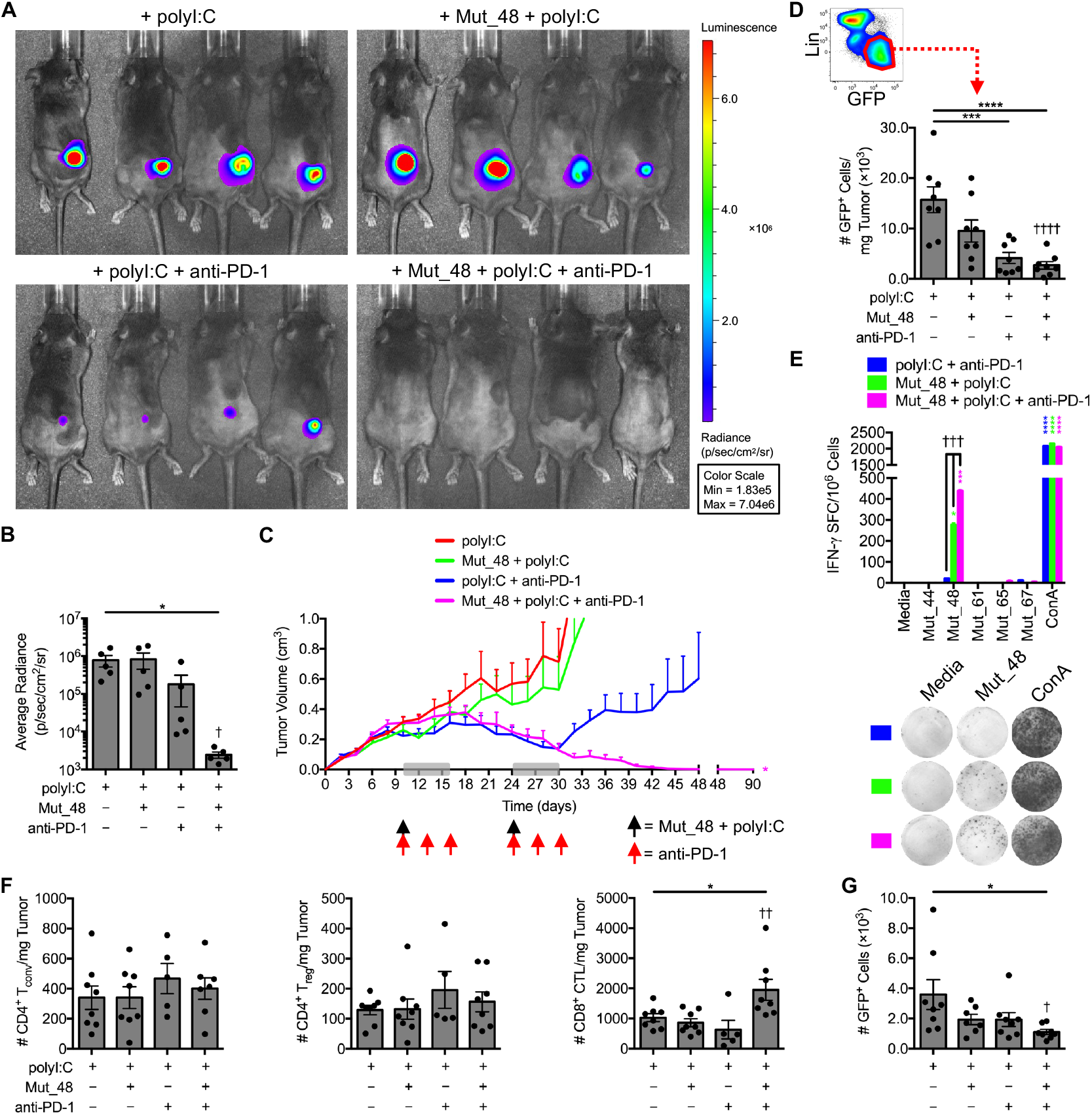
Delayed therapeutic co-delivery of anti-PD-1 and *Cltc* Δ15 promotes clearance of established SCC VII tumors. C3H/HeJ mice injected with 5×10^5^ live SCC VII-Luc/GFP cells and given 50 μg polyI:C alone or in combination with 5 μg Mut_48 peptide at day 10 post-challenge (black arrow). Select groups of mice also received anti-PD-1 at days 10, 13, and 16 (red arrow). The immunotherapy cycle repeated at day 24 (grey box). (**A** and **B**) Bioluminescence of mice at 35 days after challenge and (**C**) tumor volume kinetics tracked to day 90 (n = 6 per group). (**D**) Tumors harvested at day 17 after the first round of immunotherapy assessed for the presence of Lin^−^GFP^+^ SCC VII cells where Lin (lineage) comprised a dump gate of anti-CD31, anti-CD45, and anti-LYVE1 (n = 8 per group). (**E**) Mononuclear cells harvested from the spleens and Ig LNs of surviving C3H/HeJ mice at day 90 post-live cell challenge and assessed for IFN-γ production via ELISPOT after re-stimulation with NeoAg-pulsed BMDCs (n = 3 per group). (**F**) TIL isolated from day 17 tumor-bearing mice assessed for CD4^+^CD25^±^FoxP3^−^ T_conv_, CD4^+^CD25^+^FoxP3^+^ T_reg_, and CD8^+^ CTL (n = 5-8 per group). (**G**) Number of total Lin^−^GFP^+^ SCC VII cells in the ipsilateral Ig LN from day 17 tumor-bearing mice given therapy beginning at day 10 as a single cycle (n = 8 per group). All experiments were performed two or more times and data indicate means ± s.e.m.; (**B**, **D**, **F**, and **G**) \**P* < 0.05, \*\*\**P* < 0.001, and \*\*\*\**P* < 0.0001 (Student’s t test); ^†^*P* < 0.05, ^††^*P* < 0.01, and ^††††^*P* < 0.0001 (one-way ANOVA and Dunnett’s post hoc test relative to polyI:C); (**C**) \**P* < 0.05 (two-way ANOVA and Dunnett’s post hoc test relative to polyI:C); (**E**) \**P* < 0.05, \*\*\**P* < 0.001, and \*\*\*\**P* < 0.0001 (Student’s t test of data with SI > 2 and Poisson < 5%); ^†††^*P* < 0.01 (one-way ANOVA and Dunnett’s post hoc test relative to polyI:C + anti-PD-1).

### Neoantigen vaccination increases the presence of stem-like and intermediate exhausted CD8^+^ T cells

It is well established that CD8^+^ TIL which co-express high levels of inhibitory receptors (including PD-1 and Tim-3) exist in a terminally differentiated, exhausted state (T_ex-term_) in human cancer patients and murine tumor models (57, 58). The CD8^+^ T_ex_ lineage has a transcriptional profile and epigenetic landscape distinct from that of memory (T_mem_) and effector (T_eff_) subsets that involves rewiring via key transcription factors including TCF-1 (reinforcing stemness or memory-like features) and TOX (enforcing terminal exhaustion) (59, 60). Enrichment of PD-1^hi^Tim-3^+^TOX^+^TCF-1^−^CD8^+^ T_ex-term_ populations in tumor biopsies is directly correlated with a poor prognosis for durable responses to ICB (61). In contrast, stem-like precursor/progenitor PD-1^lo^Tim-3^−^TOX^+/−^TCF-1^+^CD8^+^ T cells (T_prec/prog_) located in tumors and/or peripheral lymphoid organs specifically expand in response to PD-(L)1-based ICB and differentiate into PD-1^lo^Tim-3^+^ effector-like cells marked by expression of the chemokine receptor, CX3CR1 (14, 57, 59, 62–65). Intermediate CX3CR1^+^CD8^+^ T_ex_ (T_ex-int_) are transitory between stem-like T_prec/prog_ and T_ex-term_ states, can be cytotoxic and produce granzyme B (GzmB) and IFN-γ, may resemble short-lived T_eff_ arising after acute antigen exposure as both express KLRG-1, and yet are distinguished from short-lived T_eff_ by TOX expression (59). In all cases, ICB does not prevent eventual terminal exhaustion as all CD8^+^ T_ex_ subsets (T_prec/prog_ > T_ex-int_ > T_ex-term_) are epigenetically scarred shortly after priming, with none being able to form T_mem_ (66–68). Moreover, it is speculated that lack of CD4^+^ T cell help during CD8^+^ T cell priming (known to drive durable T_mem_ formation) is linked with acceleration of T_ex_ differentiation as helpless CD8^+^ T cells and CD8^+^ T_ex-term_ transcriptionally resemble one another (23, 69).

We therefore sought to examine how Mut_48 vaccination and anti-PD-1 treatment reshape the CD4^+^ and CD8^+^ T cell landscape across the TME and periphery. To this end, SCC VII tumor-bearing mice were treated with SC Mut_48 + polyI:C mixtures and/or IP anti-PD-1 at day 10 following tumor inoculation, and CD45^+^ cells were purified from TIL, splenocyte, and tumor-draining Ig LN fractions at the day 17 effector phase and processed for high-dimensional fluorescence-activated cell sorting (FACS). Gated T cells were concatenated from all organs and projected into uniform manifold approximation and projection (UMAP) space using 21 phenotypic features known to define naïve CD4^+^/CD8^+^ T cells (T_n_), CD4/8^+^ T_eff/mem_, CD4^+^ T_reg_, CD8^+^ T_prec/prog_, CD8^+^ T_ex-int_, and CD8^+^ T_ex-term_ subsets. Clear separation of total CD4^+^ and CD8^+^ T cells was achieved (Figure 6A). CD8^+^ T cell subpopulations from TIL, spleen, and LN appeared entirely distinct, whereas CD4^+^ T cells from TIL and LN appeared to occupy a similar space separate from the spleen (Figure 6B, upper panel). When viewing TIL positioning alone in relation to treatment, a strong association between Mut_48 vaccination and disappearance of select CD4^+^ and CD8^+^ T cell subpopulations was observed, whereas anti-PD-1 treatment either caused more subtle shifts or appeared to expand pre-existing subpopulations relative to polyI:C treatment alone (Figure 6B, lower panel).

**Figure 6.**
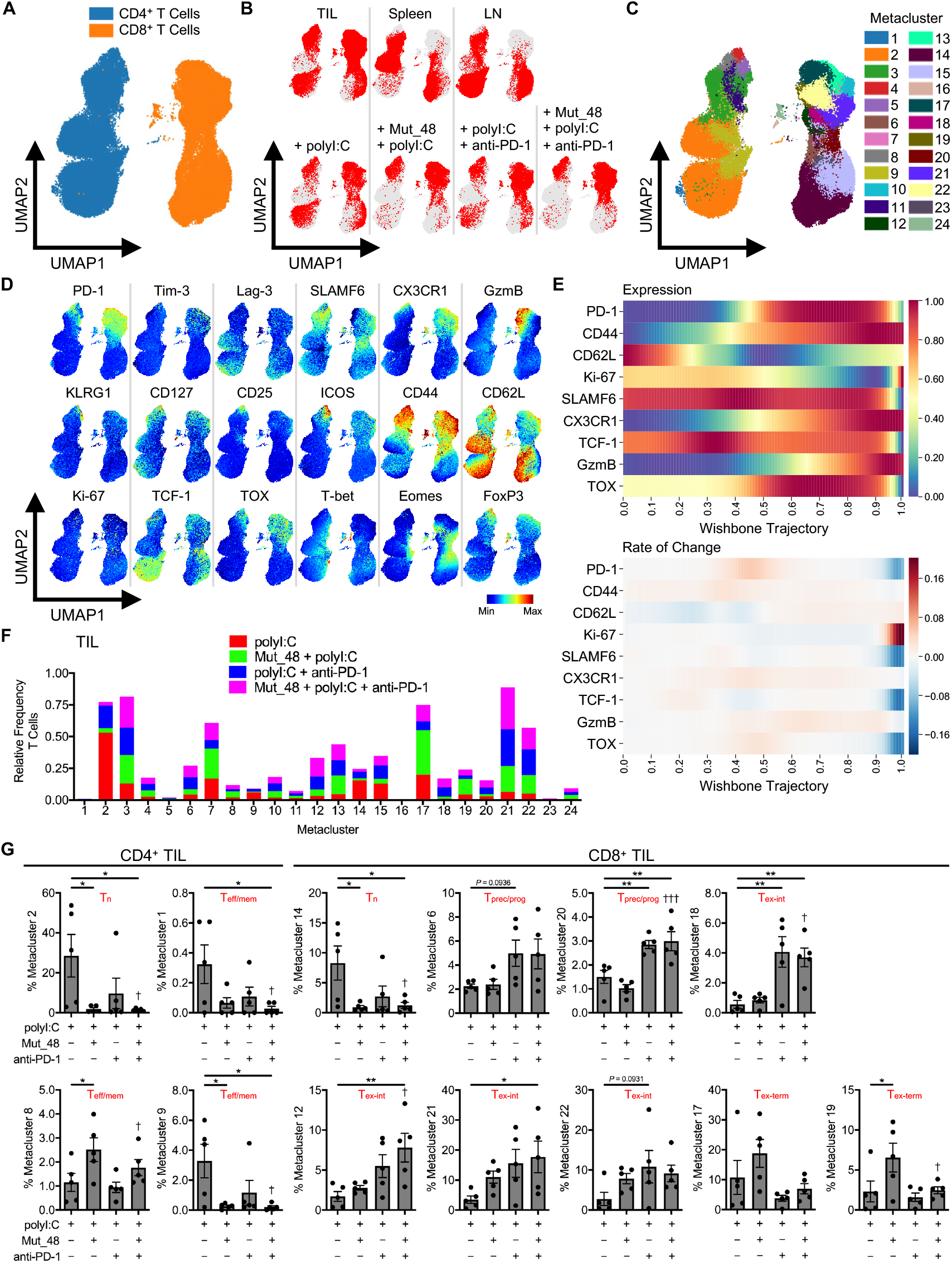
*Cltc* Δ15 vaccination enhances priming and refocuses the anti-PD-1-induced CD8^+^ T cell response towards intermediate, effector-like subsets in tumors. C3H/HeJ mice injected with 5×10^5^ live SCC VII-Luc/GFP cells and given 50 μg polyI:C alone or in combination with 5 μg Mut_48 peptide at day 10 post-challenge. Select groups of mice also received anti-PD-1 at days 10, 13, and 16. TIL, tumor-draining Ig LN, and spleens isolated from day 17 tumor-bearing mice gated on total CD4^+^ and CD8^+^ T cells (n = 5 per group). (**A**) High-dimensional FACS UMAP of all organs and treatments colored by CD4^+^ versus CD8^+^ T cell type with (**B**) total T cell positioning annotated by organ (upper panel) and TIL alone annotated by treatment (lower panel). (**C**) Identified T cell metaclusters and (**D**) expression profiles of selected phenotypic markers associated with each metacluster in UMAP space. (**E**) Pseudotime trajectory of CD8^+^ T cells using Wishbone analysis with CD8^+^ T_n_ (0.0 = start) to T_ex-term_ and short-lived T_eff_ (1.0 = end) differentiation displayed and branches in development converged. Distribution of CD62L (T_n_ and T_mem_), TCF-1/SLAMF6 (T_prec/prog_ and T_mem_), GzmB/CX3CR1/CD44/Ki-67 (T_ex-int_ and T_eff_), and TOX/PD-1 (T_ex-int_ and T_ex-term_) represented as expression (upper panel) and rate of change (lower panel). (**F**) Stacked bar plot of treatment type distribution across T cell metaclusters in TIL. (**G**) Frequency of treatment type contribution to significant and selected CD4^+^ TIL (left panel) and CD8^+^ TIL (right panel) metaclusters of interest from (**F**) with assigned cell type displayed based on (**C** and **D**). All experiments were performed two or more times and data indicate means ± s.e.m.; (**G**) \**P* < 0.05 and \*\**P* < 0.01 (Student’s t test); ^†^*P* < 0.05 and ^†††^*P* < 0.001 (one-way ANOVA and Dunnett’s post hoc test relative to polyI:C).

To gain a more detailed perspective on how treatment affected T cell differentiation, 24 metaclusters were identified that captured the granularity observed in marker profiles across the UMAP field (Figure 6, C and D). This map revealed that treatments across organs had captured the CD8^+^ T_ex_, T_eff_, and T_mem_ lineages, where pseudotime trajectory analysis of gated CD8^+^ T cells showed highly coordinated expression patterns between CD62L (T_n_ and T_mem_), TCF-1/SLAMF6 (T_prec/prog_ and T_mem_), GzmB/CX3CR1/CD44/Ki-67 (T_ex-int_ and T_eff_), and TOX/PD-1 (T_ex-int_ and T_ex-term_) (Figure 6E). Taken together, CD4^+^/CD8^+^ T_n_ (metaclusters 2 and 14), CD4^+^/CD8^+^ T_eff/mem_ (metaclusters 1, 3, 8, 9, 11, 15, 16, 23, and 24), CD4^+^ T_reg_ (metaclusters 4 and 5), CD8^+^ T_prec/prog_ (metaclusters 6 and 20), CD8^+^ T_ex-int_ (metaclusters 7, 10, 12, 13, 18, 21, and 22), and CD8^+^ T_ex-term_ (metaclusters 17 and 19) were identified based on these criteria (Figure 6, C-E).

Metaclusters were next parsed by frequency of CD4^+^ T cells or CD8^+^ T cells among treatments in TIL (Figure 6, F and G), spleen (Figure 7, A and B), and tumor-draining Ig LN (Figure 7, C and D). Of the 24 metaclusters, we found four CD4^+^ T cell- and six CD8^+^ T cell-associated metaclusters to display statistical significance relative to polyI:C treatment alone in TIL. Within CD4^+^ TIL, we observed a significant decrease in T_n_ and T_eff/mem_ cells (metaclusters 1, 2, and 9) associated with Mut_48 vaccination consistent with T cell priming. We also observed that Mut_48 vaccination caused an increase in CD4^+^ T_eff/mem_ metacluster 8, which appeared to be of possible T follicular helper cell (T_fh_) origin based on heightened PD-1 and ICOS expression (Figure 6G and Figure 7E left panels). Across all organs and treatments, CD4^+^ T_eff/mem_ (non-T_reg_) did not display markers of cytotoxicity (KLRG-1 and GzmB) consistent with their role as helpers within this model (Figure 6D and Figure 7E). Within CD8^+^ TIL, we also observed a decrease in T_n_ (metacluster 14) associated with Mut_48 vaccine induced priming. Anti-PD-1 treatment appeared to be sufficient to cause the expansion of T_prec/prog_ subsets (metaclusters 6 and 20). Select T_ex-int_ subpopulations could be supported by anti-PD-1 alone (metacluster 18), Mut_48 and anti-PD-1 combination (metaclusters 12 and 21), or either treatment (metacluster 22). T_ex-term_ subpopulations (metaclusters 17 and 19) were observed to be expanded by Mut_48 vaccination alone; however, this was halted by anti-PD-1 treatment, known to mobilize PD-1^−^ T_n_ and PD-1^lo^ T_prec/prog_ at the expense of PD-1^hi^ T_ex-term_ cells (Figure 6G and Figure 7E right panels) (70, 71). While no difference was observed in Mut_48 and anti-PD-1 combination supporting T_prec/prog_ differentiation in TIL over either treatment alone, we did note that small populations of T_prec/prog_ (metacluster 20) were significantly expanded after combination treatment in both spleen and tumor-draining Ig LN (Figure 7, A-D). These data suggest that combining PD-1 blockade with NeoAg peptide vaccination leads to an outgrowth of non-cytotoxic, helper CD4^+^ T cell subsets and more effectively expands pre-exhausted CD8^+^ T_prec/prog_ in the periphery and T_ex-int_ populations in the TIL fraction.

**Figure 7.**
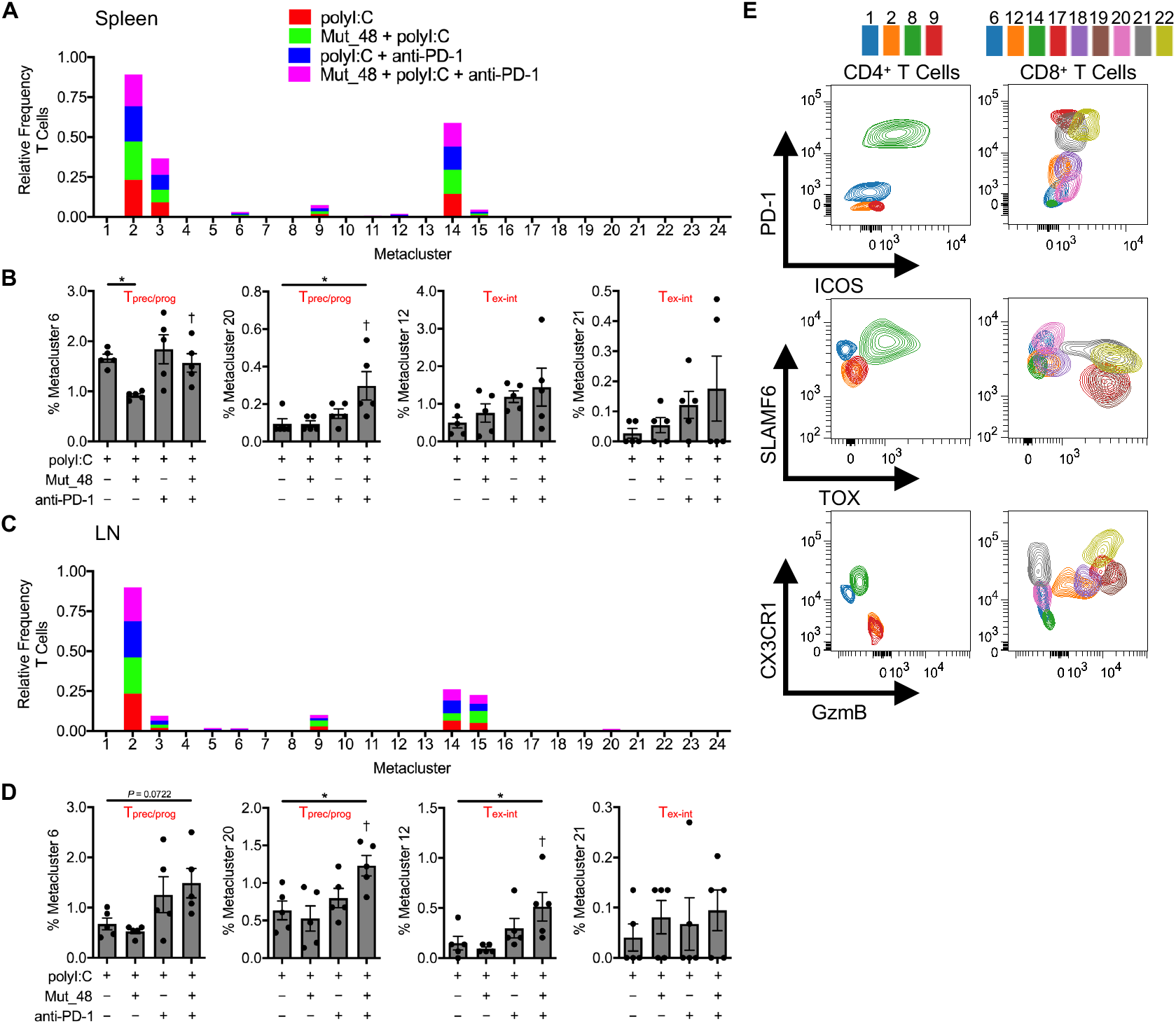
Combining anti-PD-1 and *Cltc* Δ15 expands precursor/progenitor and intermediate exhausted CD8^+^ T cells in peripheral lymphoid organs. C3H/HeJ mice injected with 5×10^5^ live SCC VII-Luc/GFP cells and given 50 μg polyI:C alone or in combination with 5 μg Mut_48 peptide at day 10 post-challenge. Select groups of mice also received anti-PD-1 at days 10, 13, and 16. (**A** and **B**) Spleens and (**C** and **D**) tumor-draining Ig LN isolated from day 17 tumor-bearing mice (n = 5 per group). Stacked bar plot of treatment type distribution across T cell metaclusters in (**A**) spleens and (**C**) tumor-draining Ig LN. Frequency of treatment type contribution to selected (**B**) spleen and (**D**) tumor-draining Ig LN T cell metaclusters of interest with assigned cell type displayed. (**E**) Representative FACS profiles of metaclusters of interest from all organs converged. All experiments were performed two or more times and data indicate means ± s.e.m.; (**B** and **D**) \**P* < 0.05 (Student’s t test); ^†^*P* < 0.05 (one-way ANOVA and Dunnett’s post hoc test relative to polyI:C).

### T_h_1-based help from CD4^+^ T cells is required for neoantigen vaccine efficacy

The observation that the Mut_48 peptide, containing both a CD4^+^ and CD8^+^ T cell epitope, was the most effective NeoAg against a tumor lacking MHC II prompted us to investigate the functional contribution made by *Cltc*-specific CD4^+^ T cells towards therapeutic vaccination. Based on our findings in TIL subset identification, linkage with treatment, and overall mechanism of action of tethered helper-effector epitopes within a single peptide, we hypothesized that NeoAg-induced mobilization of CD4^+^ T cells was completely reliant on CD40-mediated signaling, known to both directly relay T_h_1-based help via an APC and support the development of T_fh_ cells in turn sustaining IL-21-biased support of CD8^+^ CTL (23, 26, 29, 30). CD4^+^ T cell depletion before NeoAg peptide vaccination resulted in partial tumor control, suggesting that helpless CD8^+^ T cells primed in the absence of CD4^+^ T cells are not fully effective, whereas depletion of CD8^+^ T cells just prior to therapy led to rapid tumor growth, showing that CD8^+^ T cells are required as effectors against SCC VII. Agonistic anti-CD40 cross-linking antibody fully restored CD8^+^ T cell-mediated tumor rejection in the complete absence of CD4^+^ T cells (Figure 8A), suggesting that providing T_h_1-based cell help is a key feature for effective therapy even in the absence of CD4^+^ T_fh_. Consistent with this, we found that the *Cltc* CD4^+^ T cell epitope could be replaced by PADRE(X) when tethered to the CD8^+^ T cell minimal epitope (Mut_48.10) in a vaccine/PD-1 blockade therapeutic combination regimen and still result in complete tumor rejection (Figure 8B).

**Figure 8.**
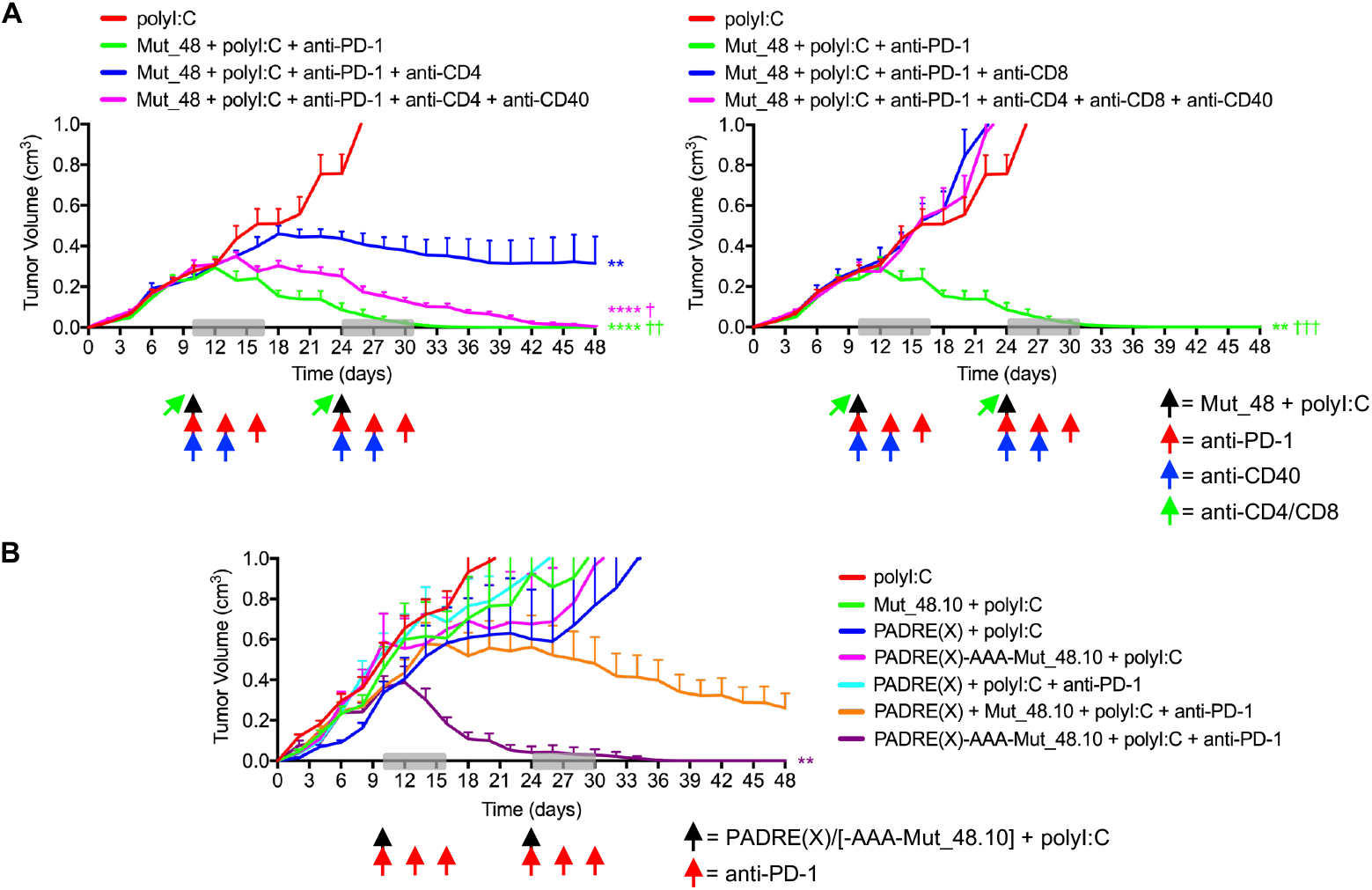
Tethered CD4^+^ T cell helper epitopes optimize checkpoint blockade- and CTL-mediated SCC VII tumor destruction via a CD40-dependent mechanism. (**A**) C3H/HeJ mice injected with 5×10^5^ live SCC VII-Luc/GFP cells and given 50 μg polyI:C alone or in combination with 5 μg full-length Mut_48 peptide at day 10 post-challenge (black arrow). Peptide-treated mice also received anti-PD-1 at days 10, 13, and 16 (red arrow). The immunotherapy cycle repeated at day 24 (grey box). CD4^+^ and CD8^+^ T cells were depleted 1 day prior to each immunotherapy cycle (green arrow), and anti-CD40 was delivered as indicated (blue arrow) with resultant tumor volume kinetics (n = 6 per group). (**B**) Tumor-bearing C3H/HeJ mice delivered 50 μg polyI:C alone or combined with 5 μg PADRE(X), Mut_48.10, mixed PADRE(X) and Mut_48.10, or tethered PADRE(X)-AAA-Mut_48.10 peptide at day 10 post-challenge (black arrow) and anti-PD-1 (red arrow) as in (**A**) with resultant tumor volume kinetics (n = 6 per group). All experiments were performed two or more times and data indicate means ± s.e.m.; (**A** and **B**) \*\**P* < 0.01 and \*\*\*\**P* < 0.0001 (two-way ANOVA and Dunnett’s post hoc test relative to polyI:C); ^†^*P* < 0.05, ^††^*P* < 0.01, and ^†††^*P* < 0.001 (two-way ANOVA and Dunnett’s post hoc test relative to Mut_48 + polyI:C + anti-PD-1 + anti-CD4 or anti-CD8 [blue groups]).

Given SCC VII-derived CD44^hi^ tCSC had increased PD-L1 expression (Supplemental Figure 3E), ICB resistance was lastly evaluated in this context after one cycle of therapeutic contralateral SC Mut_48 + polyI:C mixtures and/or IP anti-PD-1 deliveries to mice bearing day 10 SCC VII-Luc/GFP tumors. Mut_48 vaccination and PD-1 blockade individually did not result in significantly increased active caspase-3 in either SCC VII CD44^lo^ or CD44^hi^ subset. In contrast, PD-1 blockade combination with the Mut_48 NeoAg vaccine distinctly resulted in the targeting of both SCC VII CD44^lo^ and CD44^hi^ subsets (Supplemental Figure 10). These findings collectively suggest that the presence of different types of tumor resistance mechanisms (either to chemotherapy or checkpoint blockade monotherapy) can be effectively overcome by a functionally rationalized NeoAg vaccine combination approach.

## Discussion

This study advances our understanding of the therapeutic use of NeoAg in the setting of personalized cancer vaccines. First, we demonstrate that the natural T cell response to cell-associated tumor antigen can be functionally queried in an MHC agnostic manner to identify NeoAg targets for both CD4^+^ and CD8^+^ T cells. Second, by directly analyzing T cell responses primed by the intact immune system against irradiated tumor cells under physiologic conditions, only responses to natural NeoAg ligands were detected. We selected 39 mutations based on expression levels and found that four of these could be validated as NeoAg, with three recognized by CD4^+^ T cells and one by both CD4^+^ and CD8^+^ T cells, with therapeutic activity contained in the *Cltc* mutation targeted by both subsets (Mut_48). Third, our findings demonstrate that synthetically linking a NeoAg-specific or universal CD4^+^ T cell epitope with a NeoAg-specific CD8^+^ T cell epitope in a single vaccine construct allows optimal stimulation of the endogenous immune system in a tumor-bearing host to effectively mediate complete rejection of a large primary tumor burden and metastases. This could lead to novel vaccination strategies in which a pan-DR epitope such as PADRE could be tethered to validated CD8^+^ T cell targets as synthetic hybrid peptides if a suitable tumor-specific helper epitope is unavailable (54). Lastly, we show that chemotherapeutic and ICB resistance of neoplasms dominated by PD-1^hi^Tim-3^+^TOX^+^TCF-1^−^CD8^+^ T_ex-term_ and PD-L1^+^ tCSC presence in the TME are effectively overcome by strategized combination with NeoAg vaccination provided the newly recruited effector tumor-specific CD8^+^ T cells are helped by CD4^+^ T cells.

Although MHC binding prediction algorithms can reduce the number of mutations to be considered as candidate NeoAg, they cannot inform on which mutations will be naturally processed and presented at the surface of a tumor cell expressing the source protein or cross-presented by professional APC (27). In a study relying on MHC I prediction for the identification of NeoAg amongst mutations found in murine B16 melanoma, CT26 colorectal, and 4T1 mammary carcinoma models, 21-45% of filtered mutations were immunogenic and elicited T cell IFN-γ production; however, only 2-13% of these were MHC I-restricted (72). MHC II prediction is even less reliable due to the open structure of the MHC II binding groove, which accommodates peptides of varying lengths (72, 73). This is highlighted in a recent report in which a single mutation amongst 24 (4.2%) predicted for MHC II presentation was found to be a target of CD4^+^ T cells and capable of enhancing prophylactic CD8^+^ T cell immunity in a highly immunogenic sarcoma model (74). In this present work, we highlight that NeoAg filtration based on the direct ex vivo biological activity of CD4^+^ and CD8^+^ T cells provides a more rapid and accurate method for screening the function of these cells for streamlined translation in the design of NeoAg vaccines.

We speculate that NeoAg containing naturally linked or overlapping CD4^+^ and CD8^+^ T cell epitopes are rarely found within the onco-proteome, although a large-scale ex vivo effector cytokine-based functional study could demonstrate the prevalence and practical usefulness of this NeoAg subset across cancer types. Vaccination of melanoma patients in recent clinical trials using either synthetic long peptide or RNA-based vectoral approaches filtered for predicted MHC I binding revealed that anti-tumoral CD8^+^ T cell reactivity was accompanied by a significant CD4^+^ T cell MHC II-restricted component or sometimes contained single epitopes dually recognized by both CD4^+^ and CD8^+^ T cells (75, 76). A positive correlation thus appears to exist post-treatment between the presence of overlapping CD4^+^ and CD8^+^ T cell epitopes and the observation of durable responses to NeoAg therapy. Although these studies demonstrate that vaccination by predictive NeoAg filtration is generally feasible for the treatment of metastatic melanoma, it is unclear if the inclusion of epitopes yielding no IFN-γ response or those eliciting single CD4^+^ or CD8^+^ T cell responses negatively impact vaccine design via antigenic competition for MHC presentation (28).

This investigation and recent findings by Westcott *et al.* jointly demonstrate that low NeoAg expression and poor CD8^+^ T cell priming can be overcome by sustaining NeoAg vaccine-induced responses using a combination of CD40 cross-linking and ICB (8). We additionally found CD4^+^ T cell helper epitopes to be equally effective in supporting *Cltc*-specific CD8^+^ T cell responses in vaccines formulated as completely tumor-specific (contained within the same or distinct NeoAg) or tumor-nonspecific (application of a universal helper epitope), suggesting that tumor-specific endogenous effector functions of CD4^+^ T cells (Figure 1D) might be dispensable in our model during therapeutic vaccination. These data directly contrast observations made by Alspach *et al*. in a bilateral T3>KP sarcoma tumor model in which opposing MHC II^−^ tumors expressing a single CD8^+^ T cell epitope versus dual CD4^+^/CD8^+^ T cell epitopes only led to clearance of the latter tumor after ICB (74). The authors went on to demonstrate that induction of tumor-associated iNOS^+^ macrophages and optimal expansion of antigen-specific CD8^+^ T cells within the TME strictly require local expression of MHC II-restricted NeoAg during ICB (74). While SCC VII treated at a comparable early time point are sensitive to ICB (Figure 4, A and B), later stage SCC VII tumors are ICB-resistant (Figure 5C) despite expression of both MHC I- and MHC II-restricted NeoAg. Late-stage tumors may have a more established immunosuppressive TME and higher frequency of CD8^+^ T_ex-term_, which could limit local activities of CD4^+^ T cells. Furthermore, the induction of iNOS^+^ macrophages by CD4^+^ T cells in the TME was demonstrated to be dependent on antigen secretion by another study (77). Alspach *et al.* speculate that the mutated integrin subunit in their study may be similarly available to local APC populations, as it resides on the plasma membrane. In contrast, all of the MHC II-restricted NeoAg identified in this study are cytoplasmic proteins that may therefore be unable to induce local macrophage activation.

In the clinic, it is also observed that an increase in patient survival due to PD-(L)1-based ICB treatment of late stage disease strongly relies on peripheral expansion of both PD-1^−^CD8^+^ T_n_ and stem-like PD-1^lo^Tim-3^−^TOX^+/−^TCF-1^+^CD8^+^ T_prec/prog_, CXCR3/CCR5-mediated trafficking to tumors, and subsequent clonal replacement of pre-treatment ICB-refractive terminally exhausted PD-1^hi^Tim-3^+^TOX^+^TCF-1^−^CD8^+^ T_ex-term_ clonotypes (70, 78, 79). We hypothesize that tumor-specific CD8^+^ T cell responses are largely derived from the periphery in NeoAg/ICB-based immunotherapies and can remain ignorant of CD4^+^ T cell specificity in a vaccine, as long as help is provided. However, we do not discount that local interactions between CD4^+^/CD8^+^ T cells and APC occur in the TME, which may further explain the results by Alspach *et al*. Even in the presence of massive peripherally derived clonal replacement, small populations of preexistent CD8^+^ TIL are observed to expand locally within tumors after ICB (70). These local responses may be attributed to recently discovered CD8^+^ T cell:APC interactions directly in the TME (80). Outside of CD4^+^ T_h_1 CD40-mediated help within peripheral LN APC populations such as conventional type 1 dendritic cells (cDC1), CD4^+^ T_fh_ may also locally provide IL-21 to CD8^+^ T cells in the TME or within tertiary lymphoid structures to support nearby expansion and effector activity (26, 29, 30, 81). We found that NeoAg and ICB combination enhanced the amount of intratumoral PD-1^hi^ICOS^+^CD4^+^ T_fh_-like cells (Figure 6G and Figure 7E left panels); however, agonistic cross-linking of CD40 overcame the lack of CD4^+^ T cells (both T_h_1 and T_fh_), suggesting that targeting peripheral APC, including cDC1, for relay of help is necessary and sufficient to form a stable anti-tumor CD8^+^ T cell response. Although cross-linking CD40 did achieve complete tumor regression in the absence of CD4^+^ T cells, it did occur at a slower kinetic compared to tumor-bearing animals having an intact CD4^+^ T cell population (Figure 8A). Therefore, in addition to CD4^+^/CD8^+^ T cell NeoAg vaccination programming help at priming in the tumor-draining LN, it may also induce other CD4^+^ T cell responses at later time points, such as the arrival of CD4^+^ T_fh_-like cells in the TME as we highlight in this work, to improve the quality of the CD8^+^ T cell response—as enhanced effector function and/or ability to resist exhaustion. Biomarkers that define when CD4^+^ T cells can directly reshape the TME and if local APC populations can relay CD4^+^ T_h_1-versus T_fh_-biased cell help to CD8^+^ T_n_, T_eff_, T_prec/prog_, and/or T_ex-int_ remain to be elucidated.

While persistent antigen is thought to predominantly lead to the formation of CD8^+^ T_ex_ in the TME, it is speculated that lack of CD4^+^ T cell help and/or deficient accessory co-stimulation by members of the immunoglobulin and tumor-necrosis factor receptor (TNFR) superfamilies during priming also play a role in initiating and/or accelerating the exhaustion program (18, 23, 69). In SCC VII where PD-1^hi^Tim-3^+^TOX^+^TCF-1^−^CD8^+^ T_ex-term_ dominate the TME, we show that application of the Mut_48 NeoAg vaccination alone in a therapeutic setting did not cause a shift in this population towards T_ex-int_ or T_eff_ phenotypes, suggesting that late provision of CD4^+^ T help and CD8^+^ T cell priming is not sufficient in this window nor is of adequate magnitude to prevent exhaustion of the *Cltc*-specific response. Blockade of PD-1 is instead necessary to reinvigorate the response, upon which NeoAg co-administration acts to enhance upstream CD8^+^ T_prec/prog_ presence in the periphery and downstream CX3CR1^+^GzmB^lo/−^CD8^+^ T_ex-int_ numbers in the TME. Vaccine-mediated reshaping of the CD8^+^ T cell response was better able to support both real-time tumor rejection of resistant PD-L1^+^ tumor cell subsets and stabilize CD8^+^ T_mem_ formation. The Mut_48 NeoAg used in the design of this vaccine is of low predicted MHC I affinity (4988.7 nM IC_50_) using NetMHCpan (v4.0) (53). It has recently been shown that NeoAg density and T cell receptor (TCR) affinity can dictate CD8^+^ T_ex_ differentiation, where low density/affinity antigens favor T_prec/prog_ and T_ex-int_ development and high density/affinity accelerates short-lived T_eff_ and T_ex-term_ formation (8, 20, 65). Thus, it is likely that vaccines designed to contain an MHC I-bound NeoAg within a specific low MHC and/or TCR affinity window, when coupled with ICB, are more probable to seed T_prec/prog_ and T_ex-int_ in tumors known to be resistant to ICB alone (*i.e.*, cold tumors or tumors otherwise dominated by PD-1^hi^Tim-3^+^TOX^+^TCF-1^−^CD8^+^ T_ex-term_ such as SCC VII) (61). More studies are needed to understand if stabilizing *Cltc* Δ15 MHC I affinity by mutating anchoring residues and/or altering TCR contact residues can selectively expand CD8^+^ T cell clones and determine T_ex_ lineage commitment. Further, since the presence of a CD4^+^ T cell epitope is required for an effective anti-tumor CD8^+^ T cell response, it remains to be determined how added CD4^+^ T cell helper epitopes dually control T_prec/prog_ and T_ex-int_ programming beyond intrinsic CD8^+^ T cell TCR signaling by low avidity epitopes at the site of the APC.

Anti-PD-1 treatment alone led to an enhanced Mut_48-specific T cell response, which was further improved by addition of prophylactic NeoAg vaccination (Figure 4C). Consistent with this study, overlaying NeoAg therapy with pembrolizumab or nivolumab (anti-PD-1) treatment of human melanoma patients does not appear to negatively impact vaccine-induced NeoAg-specific responses, but rather preserves them while also supporting epitope spreading to novel CD8^+^ T cell reactivities (55, 56, 75). Thus, beyond TCR affinity, NeoAg density, and CD4^+^ T cell help shaping CD8^+^ T_ex_ versus T_mem_ differentiation, how pre-treatment frequency of T cell clones and/or interclonal competition of the ensuing response impacts the CD8^+^ T_ex_ lineage remains unexplored.

Overall, these findings suggest that a functional approach to NeoAg identification herein described may be the most effective in terms of time, resources, and validation for the clinical implementation of personalized cancer vaccines. In this first murine proof-of-concept study, the identification of functional responses was essentially determined from memory-phase T cells isolated from peripheral lymphoid organs of mice that had actively rejected a syngeneic tumor implant. Translation of this methodology to the clinic will rely on the ability to detect these responses in real-time within the peripheral blood and/or TIL fractions of cancer patients and is currently limited strategically to cancers of low to moderate TMB. Although this report closely examines how NeoAg vaccines containing functionally determined epitopes can overcome ICB resistance, isolation and tracking of *Cltc*-specific CD4^+^ and CD8^+^ T cell TCR clonotypes in ICB-treated animals is needed to further understand the spatiotemporal and mechanistic intersection of the two immunotherapies. It will be of interest to determine whether the broad concepts explored here regarding finding NeoAg within natural tumor-specific T cell responses and in the area of strategic vaccine design can be translated to clinical interventions in the setting of human cancer immunotherapy and if advances in high-throughput techniques can amend this approach to accommodate cancers of high TMB.

## Methods

### Detailed methods

Complete methods are provided in the Supplemental Methods section.

### Data and materials availability

Data presented in this manuscript are tabulated in the main paper and in the supplementary materials. Exome-Seq and RNA-Seq metadata are archived in the NCBI sequence read archive (SRA) under the BioProject accession number PRJNA515071 and gene expression omnibus (GEO) as series GSE125078, respectively.

### Statistics

Significant differences between experimental groups were calculated using the two-tailed Student’s t test or one-way/two-way ANOVA (with group comparisons ≥ 3) where noted. Data analysis was performed using Prism software (v9.5.1) (GraphPad Software, La Jolla, CA). Values of *P* < 0.05 were regarded as being statistically significant and noted as * < 0.05, ** < 0.01, *** < 0.001, and **** < 0.0001. ELISPOT data were subject to stricter criteria and considered significant if spot # ≥ 50, *P* < 0.05, stimulation index (SI) > 2, and Poisson < 5%. Each experiment in this report was replicated 2-3 times to ensure reproducibility and reach statistical power of the results.

### Study approval

Animals were maintained/bred in the La Jolla Institute for Immunology vivarium under specific pathogen-free conditions in accordance with guidelines of the Association for Assessment and Accreditation of Laboratory Animal Care International. Human biospecimens were collected by the Moores Cancer Center Biorepository and Tissue Technology shared resource from consented patients under a University of California, San Diego (San Diego, CA) Human Research Protections Program Institutional Review Board approved protocol (HRPP #090401).

## Supporting information

Supplemental Materials for Dolina et al. 2023

## Author Contributions

Research conceptualization: JSD and SPS. Designed experiments: JSD, JL, and SPS. Performed the research: JSD, JL, SEB, SM, SMH, and RRT. Exome-Seq and RNA-Seq processing: ML, ALRP, and JAG. Analyzed data: JSD, JL, SM, EEWC, BP, and SPS. Statistical analysis: JSD, ALRP, and JAG. Writing: JSD and SPS.

## Acknowledgments

We thank the Moores Cancer Center Biorepository and Tissue Technology shared resource for biospecimen collection. We also acknowledge D. Hinz, C. Kim, W.B. Kiossis, Z. Mikulski, and R. Simmons at the La Jolla Institute for Immunology Imaging Facility for technical assistance with FACS-sorting and microscopy experiments. We additionally would like to recognize K. Jepsen and the University of California San Diego Institute for Genomic Medicine for performing Exome-Seq and RNA-Seq of tumor and control samples. M. Croft kindly assisted in critically analyzing and discussing the data. NIH grant U01 DE028227 specifically supported this publication (S.P.S. *et al.*). The Moores Cancer Center Biorepository and Tissue Technology shared resource is supported by CCSG grant P30CA23100. NIH funded equipment was supported by grants S10 RR027366 and S10 OD016262 and included BD FACSAria II cell sorters and the Illumina HiSeq 2500 System, respectively.

