## Supplemental Materials for Dolina et al. 2023 for "Linked CD4^+^/CD8^+^ T cell neoantigen vaccination overcomes immune checkpoint blockade resistance and enables tumor regression"


**This PDF file includes**


Supplemental Methods

Supplemental Figures 1-11

Supplemental Tables 1-3

Supplemental References

### Supplemental Methods

*Animals.* Female C3H/HeJ mice (The Jackson Laboratory, Bar Harbor, ME) were used in these experiments. Animals used were 8-12 weeks of age.

*Cell culturing.* The squamous cell carcinoma VII San Francisco line (SCC VII) spontaneously arose from the abdominal wall of a C3H mouse in the laboratory of Herman Suit (Harvard University, Cambridge, MA) and was adapted for partial in vitro growth by Karen K. Fu and Kitty N. Lam (University of California, San Francisco, CA). SCC VII was maintained for a maximum of 3 passages in vitro in RPMI 1640 medium containing 12.5% fetal bovine serum in all experiments except in comparisons of passage 1 to passage 8 cells. To regenerate SCC VII $P_0$ cells, C3H/HeJ mice were subcutaneously (SC) injected with $5\times10^5$ cells in 1× HBSS, tumors were harvested at day 14 after inoculation and dissociated with the mouse Tumor Dissociation kit under the tough tumor protocol (Miltenyi Biotec, Auburn, CA) followed by passage through a 70 µm cell strainer (Fisher Scientific, Pittsburgh, PA), and homogenized cells were re-seeded in vitro. For generation of SCC VII expressing luciferase and copepod-derived GFP (SCC VII-Luc/GFP), cells were transduced with the BLIV713VA-1 HIV lentiviral vector (System Biosciences, Palo Alto, CA) under 10 µg/mL puromycin selection and further purified using $GFP^+$ fluorescence-activated cell sorting (FACS).

Human tumor-initiating cancer stem cells (tCSC) isolated from head and neck squamous cell carcinoma (HNSCC) patients were generated as previously described (1). Briefly, homogenized cells from primary tumor specimens were placed over a monolayer of irradiated murine 3T3 cells in pre-conditioned 3T3 cell media containing 10 µM Y-27632 ROCK1/2 inhibitor (Enzo Life Sciences, Farmingdale, NY). After multiple passages, tCSC were weaned from 3T3 cells by differential trypsinization and maintained in media containing Y-27632 for independent cell line growth.

For experiments conducted in vitro comparing stem cells to differentiated cells, ‘basal media’ refers to RPMI 1640 medium containing 12.5% fetal bovine serum or DMEM medium containing 10% fetal bovine serum for SCC VII and HNSCC, respectively. ‘Conditioned media’ refers to media supplemented with 10 μM Y-27632.

*Tumor- and peptide-based immunizations and vaccinations.* Tumor cell-based immunization experiments in C3H/HeJ mice were conducted via SC injection of $1\times10^7$ 50 Gy irradiated SCC VII cells in 1× HBSS. 14 days later, mice were SC challenged with $5\times10^5$ live SCC VII-Luc/GFP cells in 1× HBSS ipsilaterally. For NeoAg peptide-based vaccination experiments, mice were delivered 5 μg single 20-mer peptides, 5 μg single minimal peptide epitopes, or 25 μg mixtures of 5× unique 20-mer peptides (5 μg each). Some immunization cohorts involved injection of 5 μg tethered peptides with covalently linked $CD4^+$ T cell helper and $CD8^+$ T cell minimal epitopes via a triple alanine repeat: Mut_44-AAA-Mut_48.10 and PADRE(X)-AAA-Mut_48.10. PADRE(X) contains the sequence, AKXVAAWTLKAAA where X = cyclohexylalanine, as previously described (2) but without flanking D-enantiomers as to not disrupt intracellular processing and presentation in fused epitopes. At day 21 after initial peptide injection, some experimental groups received a repeat booster regimen in the same scheme, while other groups remained untreated at this time point. All mice in peptide vaccination trials were SC challenged with $5\times10^5$ live SCC VII-Luc/GFP cells in 1× HBSS contralaterally at day 31. In both tumor and peptide immunizations, select groups included 50 μg high molecular weight polyI:C (InvivoGen, San Diego, CA) as an adjuvant at the injection site. For purely therapeutic experiments, mice bearing untreated day 10 ~300-400 $mm^3$ SCC VII-Luc/GFP tumors were given 5 μg Mut_48 20-mer or

PADRE(X)-AAA-Mut_48.10 26-mer peptides in combination with 50 μg high molecular weight polyI:C SC contralaterally and followed up with repeat injections every 14 days during the complete time course of immunotherapy trials. Live SCC VII-Luc/GFP tumors were measured by caliper every other day and by whole-animal imaging luciferase activity every 7 days in all experiments.

*Chemotherapy.* *Cis*-Diamineplatinum(II) dichloride (cisplatin; Sigma-Aldrich, St. Louis, MO) was dissolved in 0.9% sterile saline overnight. C3H/HeJ mice approximately 20 g in mass and bearing untreated day 10 ~300-400 $mm^3$ SCC VII-Luc/GFP tumors were given a single maximum tolerated dose of 6 mg/kg cisplatin in 0.9% sterile saline intraperitoneally (IP) according to previous work (3). Tumors were harvested at day 7 for flow cytometric analysis. The mass of all animals was monitored daily and with no significant changes reported during the duration of experiments.

*Antibody-mediated checkpoint blockade, $CD4^+$/$CD8^+$ T cell depletion, and assessment of help-dependence.* The GK1.5 hybridoma (ATCC, Manassas, VA) and FGK45 hybridoma originally obtained from Antonius G. Rolink (Basel Institute for Immunology, Basel, Switzerland) were cultured using in-house CELLine 1000 bioreactors (Wheaton, Millville, NJ) to obtain large quantities of anti-CD4 and anti-CD40 antibody, respectively. Anti-CD8 (116-13.1), anti-PD-1 (RMP1-14), and anti-CTLA-4 (9H10) were otherwise externally purchased for experiments (BioXCell, West Lebanon, NH). For early $CD4^+$ or $CD8^+$ T cell depletion in vaccination groups, anti-CD4 or anti-CD8 was respectively delivered IP in the following regimen: 200 μg at day −5, 100 μg at day −2, and 100 μg at day 5 relative to a day 0 immunization. For late T cell depletion after prophylactic immunization, 400 μg anti-CD4 or anti-CD8 was delivered IP 1 day prior to live tumor cell challenge. For vaccine trials, checkpoint blockade regimens began as IP injections at day 3 after live SCC VII-Luc/GFP tumor challenge with re-delivery every 3 days for a total of 5 injections. Anti-PD-1 was used at 250 µg per injection, and anti-CTLA-4 was used at 200 µg for the first injection and 100 µg for remaining injections. For peptide-based immunotherapy trials, $CD4^+$ or $CD8^+$ T cells were depleted with 400 µg anti-CD4 or anti-CD8 IP one day prior to peptide and checkpoint blockade regimens. 250 µg anti-PD-1 was then co-delivered IP alongside SC peptide and polyI:C injections and re-delivered alone twice more every 3 days following each peptide and polyI:C injection. For assessment of help-dependence in the therapeutic setting, 100 µg anti-CD40 was delivered twice intravenously (IV) just prior to and 24 hours after initiation of each round of therapy. All mice were monitored for effective T cell depletion throughout vaccine and therapeutic approaches via weekly phenotyping of peripheral blood mononuclear cells (PBMCs).

*Tumor, Spleen, and Ig LN mononuclear cell isolation.* Spleens and inguinal lymph nodes (Ig LN) were excised from the abdominal cavity of naïve and immunized mice and homogenized through sterile 70 µm cell strainers (Fisher Scientific) in RPMI 1640 medium containing 10% fetal bovine serum for all downstream applications. Tumor infiltrating lymphocytes (TIL) were collected using the mouse Tumor Dissociation kit under the tough tumor protocol (Miltenyi Biotec) prior to cell straining. For traditional FACS analysis of TIL, mononuclear cells were isolated by Lympholyte-M (Cedarlane, Burlington, NC) gradient centrifugation. For spectral analysis, leukocytes were positively selected from digested tumor samples using the EasySep Mouse $CD45^+$ Isolation kit (STEMCELL Technologies, Vancouver, BC, Canada). Viable cells were quantitated via Trypan Blue exclusion on a Vi-CELL XR cell viability analyzer (Beckman Coulter, Indianapolis, IN).

*Quantitative PCR (Q-PCR).* Total RNA was isolated using the Trizol method (Invitrogen, Carlsbad, CA), DNase I treated (Invitrogen), and reverse-transcribed using High Capacity RNAto-cDNA Master Mix (Applied Biosystems, Foster City, CA). Q-PCR was performed using Fast SYBR Green Master Mix (Applied Biosystems) on an AB StepOne Plus Real-Time PCR System. QuantiTect primers for *Mus musculus Rock1* and *Rock2* and *Homo sapiens Rock1*, *Rock2*, and *Hprt1* (Qiagen, Valencia, CA) and self-designed primers for *Mus musculus Hprt1* (forward, 5'-CTCCGCCGGCTTCCTCCTCA-3' and reverse, 5'-ACCTGGTTCATCATCGCTAATC-3') were used for detection.

*Western blotting.* Phospho(S19)-myosin light chain 2 (pMLC2) protein expression was assessed in homogenized SCC VII and HNSCC cells. Protein concentration was determined using the Pierce BCA Protein Assay kit (Thermo Scientific, Rochester, NY). Proteins were resolved, Western blotted, and incubated with anti-pMLC2 (S19) (pAb) (Cell Signaling Technology, Danvers, MA), anti-GAPDH (6C5) (Applied Biosystems), donkey anti-rabbit IRDye 800CW (pAb), and donkey anti-mouse IRDye 680RD (pAb) (LI-COR Biosciences, Lincoln, NE). Imaging was performed on an Odyssey 9120 Infrared Imaging System (LI-COR Biosciences).

*Enzyme-linked immunosorbent spot assay (ELISPOT).* Effector/memory phase mononuclear cells were isolated from spleens and Ig LNs. In select experiments, $CD4^+$ T cells were purified using the EasySep Mouse CD4 Positive Selection kit II, and $CD8^+$ T cells were negatively selected from the supernatant fraction using the EasySep Mouse $CD8^+$ T cell Isolation kit (STEMCELL Technologies). Bulk leukocytes or purified T cells were placed in co-culture with day 7 rIL-4 and rGM-CSF (Tonbo Biosciences, San Diego, CA) matured bone marrow-derived dendritic cells (BMDC) at a 10:1 ratio ($2\times10^5$:$2\times10^4$ mononuclear cells:BMDCs) in triplicate to multiscreen-IP filter 96-well plates (MilliporeSigma, St. Louis, MO) coated overnight with 2 μg/mL anti-IFN-γ (AN18) capture antibody (MabTech, Cincinnati, OH). Cells were incubated for 20 hours at 37°C, 5% $CO_2$ with 5 μg/mL test peptides or positive controls including $2\times10^4$ 150 Gy-irradiated/IFN-γ-treated and washed SCC VII cells or 5 μg/mL ConA (Sigma-Aldrich). For MHC I and II blocking, 20 μg/mL of anti-H-2K$^k$ (16-3-22S) (Novus Biologicals, Centennial, CO), anti-I-A$^k$ (10-3.6.2), or anti-I-E$^k$ (14-4-4S) (BioXCell) were added to the appropriate wells. Plates were washed with 1× PBS/0.05% Tween 20, and biotinylated anti-IFN-γ (R4-6A2) antibody (MabTech) was used for detection for 2 hours at 37°C. Development of spots proceeded via incubation of plates with Vectastain Elite ABC kit avidin-peroxidase complex (Vector Laboratories, Burlingame, CA) for 1 hour at room temperature followed by submersion in a 0.45 μm-filtered acetate buffered solution containing 3-amino-9-ethylcarbazole, *N,N*-dimethylformamide, and $H_2O_2$ (Sigma-Aldrich) for 10 minutes at room temperature. IFN-$\gamma^+$ spots were quantified on a Zeiss Stemi 2000-C stereoscope with KS ELISPOT software (v4.11) (Carl Zeiss MicroImaging GmbH, Jena, Germany) using the following parameters: spot diameter (50-500 μm), hue (90.2-9.3), saturation (16.8-94.8), contrast (3.7-100.0), shape (47.5-100.0), and slope (15.9-56.7).

*Whole-animal imaging.* Mice bearing SCC VII-Luc/GFP tumors were IP injected with 3.6 mg VivoGlo Luciferin (Promega, Madison, WI) and rested for 10 minutes. Luciferase activity was measured in mice exposed to 2% isoflurane on a 37°C platform for a 45 second exposure using an IVIS Spectrum imaging system with Living Image software (v3.2) (Caliper Life Sciences, Hopkinton, MA and Xenogen Corporation, Alameda, CA).

*Flow cytometry.* Cells were stained with fluorochrome-tagged antibodies from Abcam (Cambridge, MA), BD Biosciences (Franklin Lakes, NJ), BioLegend (San Diego, CA), Cell Signaling Technology, eBioscience (San Diego, CA), Invitrogen, Rockland Immunochemicals (Limerick, PA), Sino Biological (Wayne, PA), Thermo Scientific, and Tonbo Biosciences (Supplemental Table 3). Viable cells were indicated with the LIVE/DEAD Fixable Blue or Yellow Dead Cell Stain kits (Invitrogen). Antibody labeling of chemokine receptors was achieved by incubation of cells for 20 minutes at 37°C in RPMI 1640 medium containing 10% fetal bovine serum, followed by additional inclusion of relevant antibodies during downstream surface staining. Cell surface staining of mononuclear leukocytes or tumor cells was performed by specific antibody labeling for 15 minutes at 4°C in FACS buffer (1× PBS containing 2% fetal bovine serum and 0.1% $NaN_3$). Cells were fixed in BD Cytofix/Cytoperm (BD Biosciences). Intracellular proteins were measured in cells permeabilized with BD Perm/Wash (BD Biosciences). Nuclear protein staining was achieved using the FoxP3 Staining Set (eBioscience). Therapy induced apoptosis was assessed by intracellular staining with fluorescently labeled anti-cleaved caspase-3 (Cell Signaling Technologies). SCC VII cells were treated with 15 μM etoposide (Calbiochem/MilliporeSigma) for 20 hours to induce cell death and discriminate positive versus negative control gates of active caspase-3. The FACS gating strategy used in this study for identification of $CD4^+$ $T_{conv}$, $CD4^+$ $T_{reg}$, and $CD8^+$ CTL as well as the $Lin^-GFP^+$ SCC VII cell subsets is graphically displayed (Supplemental Figure 11). For high-dimensional FACS analysis, $CD45^+$ cells were purified to physically exclude $GFP^+$ cells from unmixing and subsequent analyses, and T cell subsets were assessed after applying additional $CD45^+$, $Thy1.2^+$, $TCR\gamma\delta^-$, and $NKp46^-$ cleanup gates (not depicted). Data were collected on a BD FACS Canto II (BD Immunocytometry Systems, San Jose, CA) or a Cytek Aurora (Cytek Biosciences, Fremont, CA) and analyzed using FlowJo (v9.9.6 and v10.0.6) software (Tree Star, Ashland, OR) or OMIQ software from Dotmatics (www.omiq.ai; www.dotmatics.com; dates accessed: 2023-04-18 to 2023-04-29), respectively. Cell sorting of $GFP^+$ SCC VII cells was performed using BD FACSAria II cell sorters (BD Immunocytometry Systems).

*Spectral data processing and analysis.* $CD4^+$ and $CD8^+$ T cells were gated as above with additional inclusion of $CD4^+CD8^+$ and $CD4^-CD8^-$ NOT gates. From this, total T cells were subsampled at

1,480 cells per sample, with 88,115 total T cells from all treatment conditions used for dimensionality reduction. Phenotypic markers not deployed in T cell gating (21 features) were used for clustering and uniform manifold approximation and projection (UMAP), where neighbors = 15, minimum distance = 0.4, components = 2, metric = euclidean, learning rate = 1, epochs = 200, and random seed = 5736. FlowSOM with consensus metaclustering was performed, where xdim = 10, ydim = 10, rlen = 10, metric = euclidean, $k$ = 12, 18, 24, 28, 32, and random seed = 4436. It was determined that $k = 24$ was the minimal value that fully captured distinct cell populations observed in the UMAP field. Trajectory analysis of select markers was performed on all $CD8^+$ T cells from all samples (39,911 total cells) with method = Wishbone, waypoints = 250, $k = 15$, random seed = 5602, and a predetermined naïve $CD8^+$ T cell ($T_n$) starting gate. Plots were generated from the Wishbone trajectory with 100 bins and represented as expression and derivative (rate of change).

*Confocal microscopy.* For comparison of murine SCC VII and human HNSCC tCSC versus differentiated cell morphology and cytoskeletons, cells were grown in Lab-Tek II 4-chambered slides (Nunc, Rochester, NY), fixed for 10 minutes in 1% periodate-lysine-paraformaldehyde (PLP), washed in 1× PBS, blocked with 2.4G2 solution (2.4G2 hybridoma supernatant containing anti-CD16/32, 30% chicken/donkey/horse serum, and 0.1% $NaN_3$), and sequentially stained with Hoechst 33342 (Invitrogen), anti-β-Tubulin AF488, Phalloidin AF555, and anti-Vimentin AF647 (Cell Signaling Technology). All images were acquired using a Zeiss LSM 780 confocal microscope equipped with a 63×/1.40 NA oil objective with 0.2 μm z-steps (Carl Zeiss MicroImaging GmbH). Image analysis was done using Imaris (v9.2) software (Bitplane AG, Zurich, Switzerland).

*Wound closure assay.* Murine SCC VII and human HNSCC cells were grown to monolayers in ibiTreat 96-well μ-plates (Ibidi, Madison, WI) in phenol-free basal versus conditioned media. Cells were scratched with a sterile cat whisker brush. The location of the cells was imaged using a Zeiss LSM 880 confocal microscope equipped with a 10×/0.45 NA air objective (Carl Zeiss MicroImaging GmbH) using transmitted light from the 633 nm LASER. 3-5 fields-of-view were tiled for each sample. Images were acquired every 60 seconds for 24 hours to monitor wound closure over time, and the imaging chamber was kept at 37°C, 5% $CO_2$ for the duration of experiments.

Image processing was performed using Fiji software (v2.0.0-rc-68/1.52n). A Laplacian filter (FeatureJ) was used to remove illumination gradients. Cells were detected with Trainable Weka Segmentation (4) and segmented based on a 50% probability threshold. In each movie, cell coverage was analyzed in the region from the scratch border (outlined manually in the first time point) to 60 μm into the wound area using a custom macro. The total area in that region covered by cells was calculated at the 0, 8, 16, and 24 hour time points.

*Bulk exome sequencing (Exome-Seq) and RNA sequencing (RNA-Seq).* Tumor tissue was collected from mice inoculated with SCC VII to perform Exome-Seq and RNA-Seq, and mouse tail samples were collected and subjected to Exome-Seq to be used as germline controls. Total DNA and RNA was purified using the AllPrep DNA/RNA Micro kit (Qiagen, Valencia, CA).

For Exome-Seq, sequencing libraries were prepared and captured using SureSelect All Exon V4 Library kits (Agilent Technologies, Santa Clara, CA) following the manufacturer's instructions. Briefly, 1-200 μg of genomic DNA from each sample was fragmented by Adaptive Focused Acoustics using an E220 Focused Ultrasonicator (Covaris, Woburn, MA) to produce an average fragment size of ~175 bp. Fragmented DNA was purified using Agencourt AMPure XP beads (Beckman Coulter). The quality of the fragmentation and purification was assessed with an Agilent 2200 Tapestation (Agilent Technologies). The fragment ends were repaired, and adaptors were ligated to the fragments. The resulting DNA library was amplified by 7 cycles of PCR, and 500 ng of each library was captured by solution hybridization to biotinylated RNA library baits for 24 hours at 65°C. Bound genomic DNA was purified with streptavidin coated magnetic Dynabeads (Invitrogen) and further amplified using 12 cycles of PCR to add barcode tags.

For RNA-Seq, total RNA was assessed for quality using an Agilent 2100 Tapestation (Agilent Technologies). RNA-Seq libraries were generated from 200 ng of total RNA using the TruSeq Stranded Total RNA Library Prep kit with Ribo-Zero Gold (Illumina, San Diego, CA). Manufacturer's instructions were followed with shear time modified to 5 minutes.

Both Exome- and RNA-enrichment libraries were sequenced on the Illumina HiSeq 2500 in Rapid Run mode (Illumina) generating 100×100 bp paired-end reads. An average Exome-Seq depth of 36.07 and RNA-Seq fold-coverage of 224.75 (~$200\times10^6$ reads) was achieved.

*Bioinformatic NeoAg identification.* Sequence reads from Exome-Seq of the tumor and normal tail samples were aligned to the mm10 reference genome using SpeedSeq Align (v0.0.3a). Exome variants were identified using SpeedSeq Somatic (v0.0.3a), and variants were annotated using SNPeff (v4.1). Reads from RNA-Seq of the tumor were aligned using STAR aligner (v2.4.1c). To confirm the expression of exome variants in the tumor, the aligned RNA data was compared with the reference genome to check the presence of exome variants using Freebayes software (v0.9.16).

Variants were considered for further analysis only if they met following criteria: sequencing depth at position of variant ≥ 10 in normal exome as well as in tumor exome and transcriptome, variant allele observations in normal exome = 0, variant allele observations in tumor RNA ≥ 10, variant allele frequency (VAF) in tumor RNA ≥ 20%, somatic score from SpeedSeq ≥ 0, single nucleotide polymorphism (SNP) quality score ≥ 20, and the variant was protein coding and protein changing. For single nucleotide variants, two 20-mer peptides containing the mutation at positions 6 and 15 for each variant in the reference protein sequence were generated. When the variant was near the start or the end of the protein sequence the mutation position was adjusted in the 20-mer peptide due to insufficient flanking amino acids. For frameshift variants, 20-mer peptides overlapping by 10 residues were generated with the first mutated amino acid starting at positions 6, 15, and so on until the first found stop codon in the mutated sequence was hit. Peptides used in experiments were synthesized by A & A Labs (DBA Synthetic Biomolecules, San Diego, CA). MHC I predictions of peptide binding to murine H-2K$^{k}$ were made using the Immune Epitope Database and Analysis Resource (IEDB) (www.iedb.org) (5, 6) NetMHCpan (v4.0) tool (7).

*Bioinformatic analysis match of murine to human RNA-Seq datasets.* Analysis match of murine SCC VII RNA-Seq TPM data represented with a cutoff > 20 to published online NCBI OncoGEO human tumor datasets was performed using Ingenuity Pathway Analysis software (IPA; v01-07) (Qiagen). Human cancer RNA-Seq data were only considered relevant if a disease versus normal comparison was conducted in the source study and patients received only radiation and/or chemotherapy (with exclusion of tumors from patients that received experimental drugs or other FDA-approved therapeutic regimens). TIL datasets were additionally omitted from this analysis. Pathways were represented with an absolute perturbed $Z$ score > |2|.

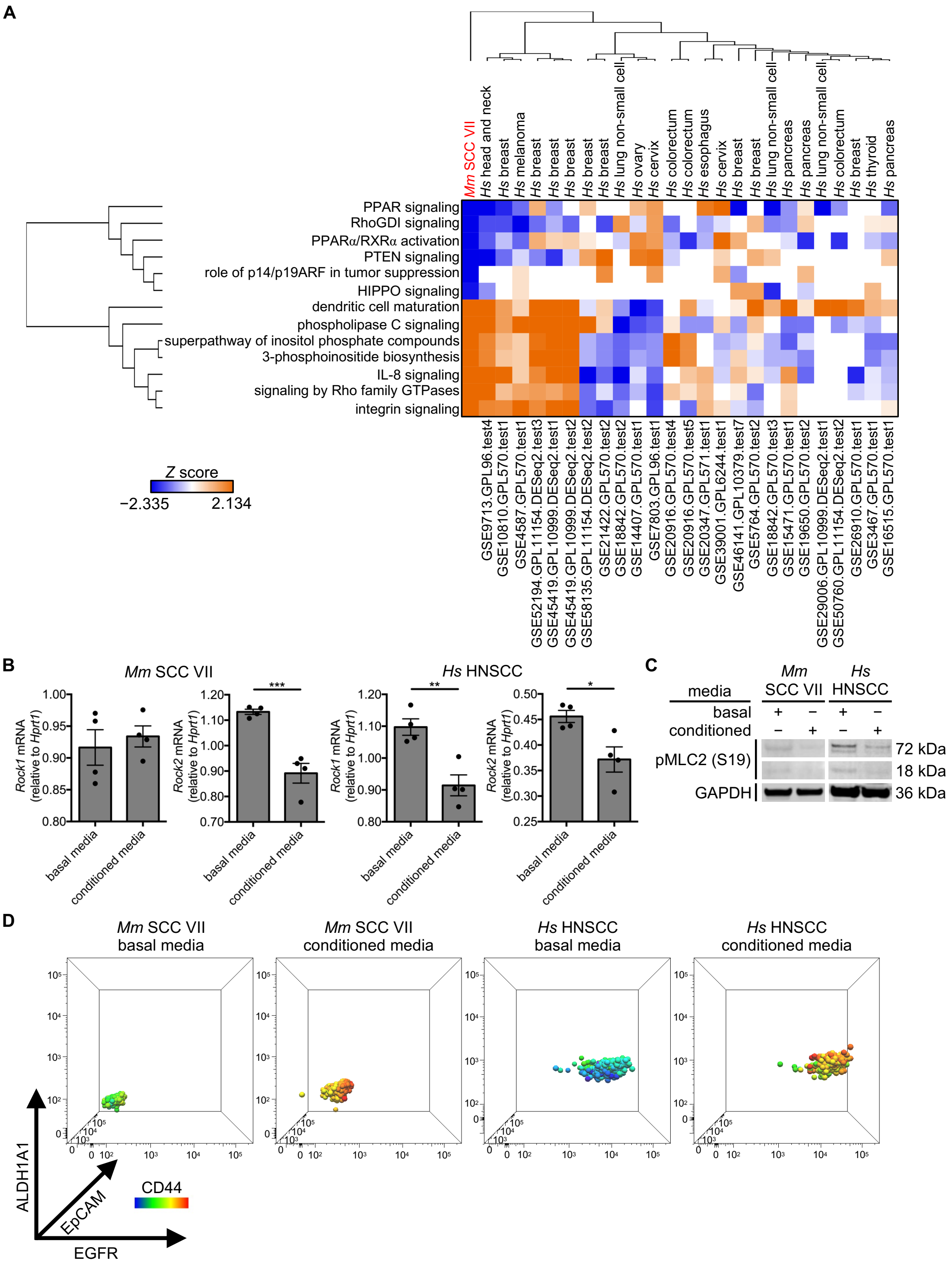


**Supplemental Figure 1. Involvement of the Rho kinase pathway in murine and human tCSC formation.** (**A**) Hierarchical clustered IPA analysis match of SCC VII RNA-Seq TPM data compared to human NCBI OncoGEO tumor datasets. Cell pathways that showed positive or negative correlation between cancers. (**B**-**D**) *Mm* SCC VII and *Hs* HNSCC cells were grown in basal media or conditioned media containing 10 μM Y-27632 ROCK1/2 inhibitor for 7 days. (**B**) Total RNA isolated from cells with *Rock1* and *Rock2* mRNA normalized to *Hprt1* mRNA quantified by Q-PCR. (**C**) pMLC2 (S19) assessed by Western blotting cell lysates. (**D**) 4D plots of surface CD44, EGFR, and EpCAM and intracellular expression of ALDH1A1 in both *Mm* SCC VII and *Hs* HNSCC (n = 4 per group). All experiments were performed two or more times and data indicate means ± s.e.m.; (**B**) *$P < 0.05$, **$P < 0.01$, and ***$P < 0.001$ (Student's t test).

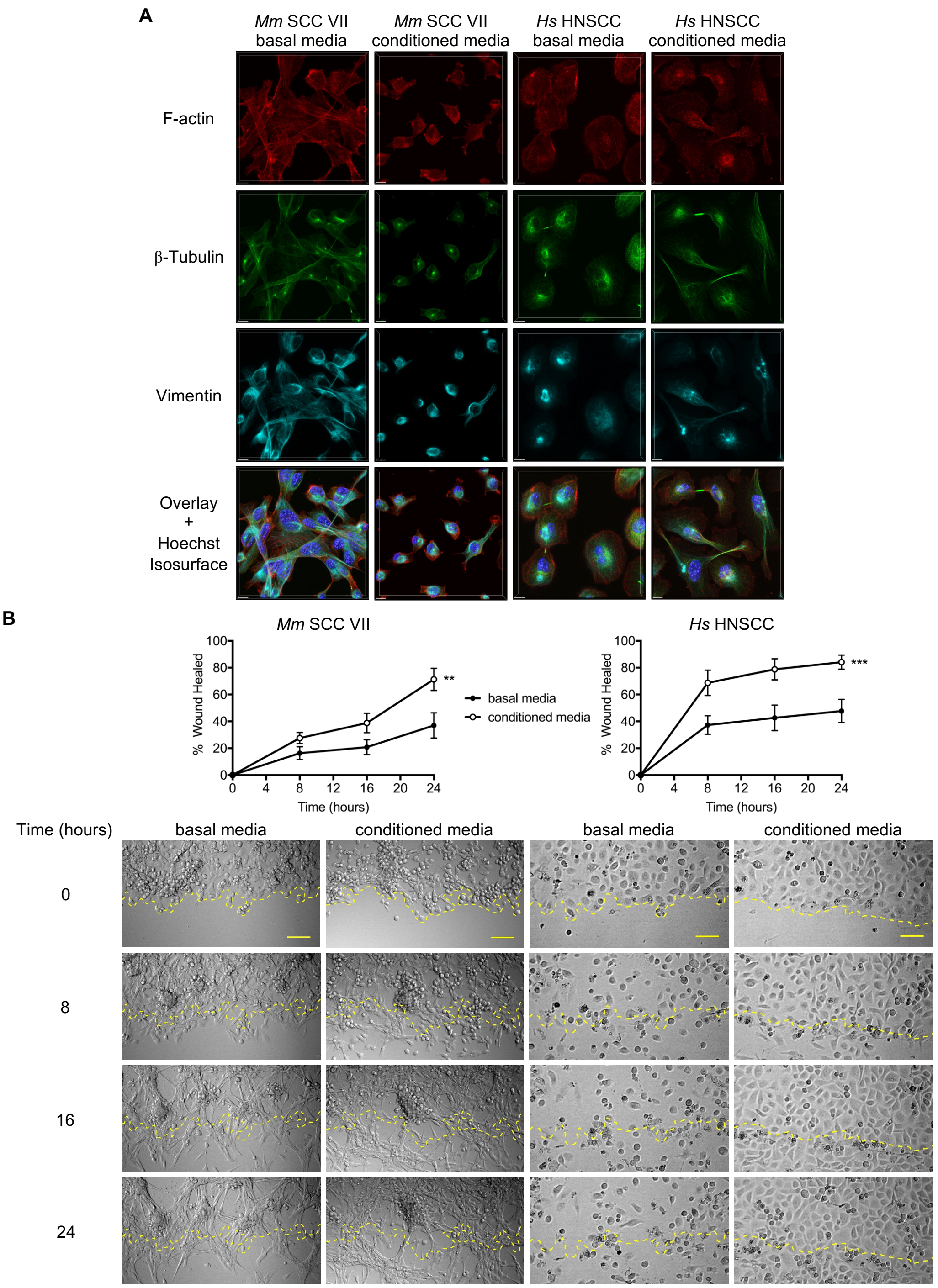


**Supplemental Figure 2. Murine SCC VII tCSC cytoskeletal disorganization and enhanced migration correlates with phenotypes observed in human tCSC.** *Mm* SCC VII and *Hs* HNSCC cells were grown in basal media or conditioned media containing 10 μM Y-27632 ROCK1/2 inhibitor for 7 days then plated for 2 additional days in chambered slides. (**A**) PLP-fixed cells stained with Phalloidin (F-actin; red), anti-β-Tubulin (green), anti-Vimentin (cyan), and Hoechst (blue isosurface in overlay images). Scale bar, 10 μm. (**B**) Wound closure assay of *Mm* SCC VII and *Hs* HNSCC cells in basal versus conditioned media over 24 hours (n = 4-6 per group). Scale bar, 100 μm. All experiments were performed two or more times and data indicate means ± s.e.m.; (**B**) $^{**}P < 0.01$ and $^{***}P < 0.001$ (two-way ANOVA and multiple comparison Sidak's post hoc test).

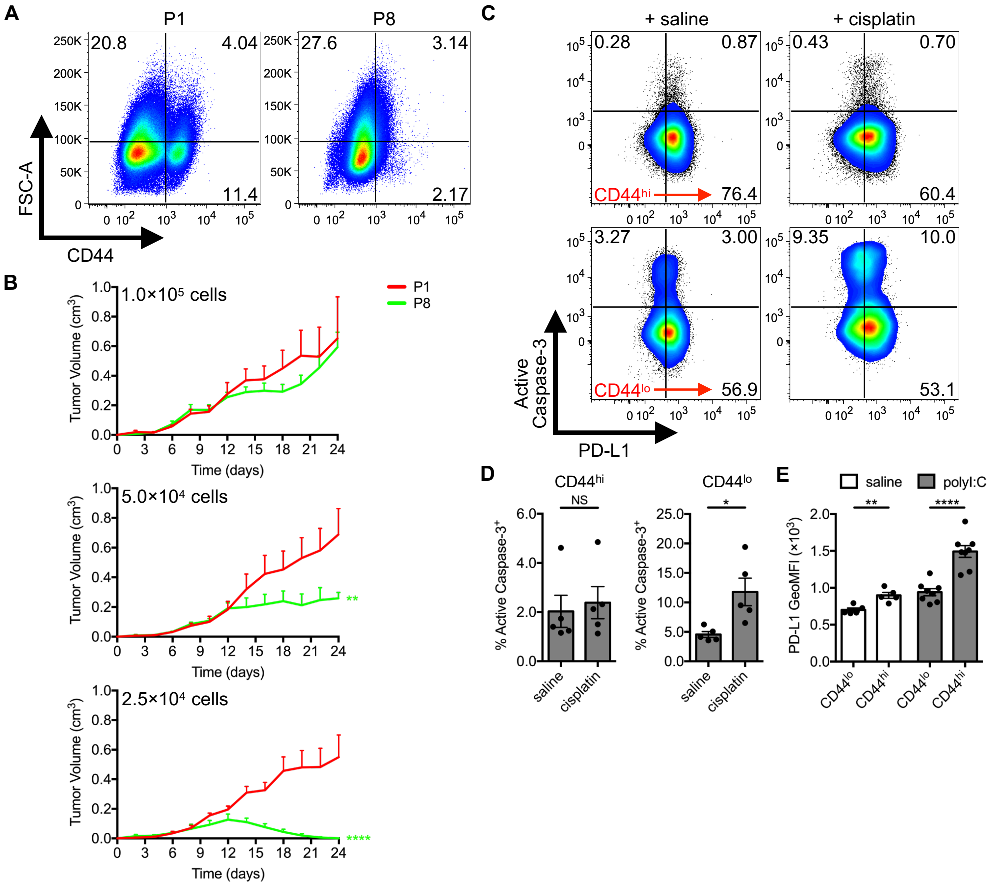


**Supplemental Figure 3. Tumorigenicity and chemoresistance of SCC VII-derived CD44$^{lo}$ differentiated cell versus CD44$^{hi}$ tCSC.** (**A**) CD44 expression of differentially passaged (P1 versus P8) SCC VII-Luc/GFP cells (n = 4 per group). (**B**) Naïve C3H/HeJ mice injected with titrations of differentially passaged live SCC VII-Luc/GFP cells with resultant tumor volume kinetics (n = 6 per group). (**C** and **D**) Tumor-bearing C3H/HeJ mice treated on day 10 with a single bolus of saline versus cisplatin with frequency of active caspase-3 in Lin$^{-}$GFP$^{+}$ SCC VII cells measured at day 17 (n = 5 per group). (**E**) PD-L1 surface expression on CD44$^{lo}$ versus CD44$^{hi}$ Lin$^{-}$GFP$^{+}$ SCC VII cells at day 17 of growth in saline- and 50 μg polyI:C-treated C3H/HeJ mice (n = 5-8 per group). All experiments were performed two or more times and data indicate means ± s.e.m.; (**B**) $^{**}P < 0.01$ and $^{****}P < 0.0001$ (two-way ANOVA and multiple comparison Sidak's post hoc test); (**D** and **E**) $^{*}P < 0.05$, $^{**}P < 0.01$, and $^{****}P < 0.0001$ (Student's t test).

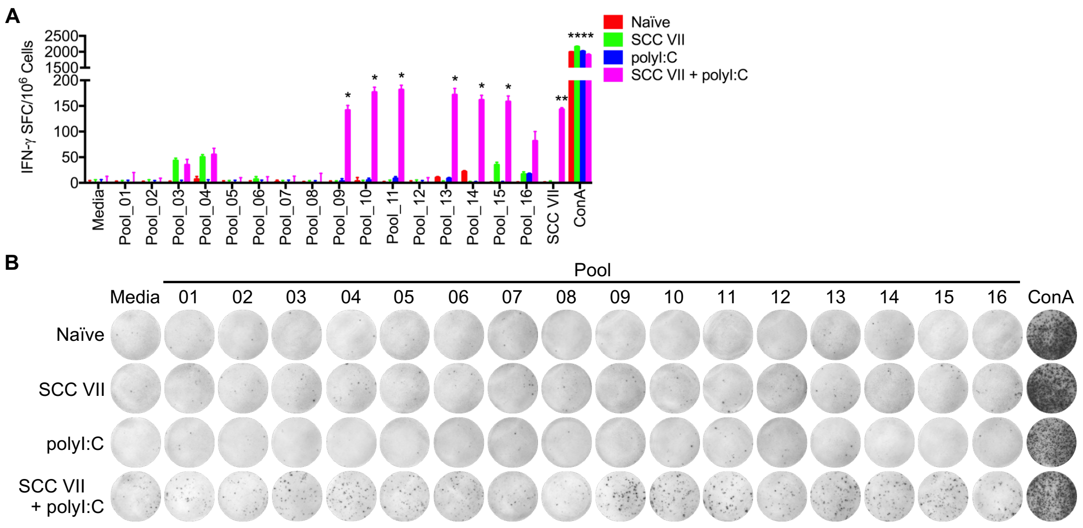


**Supplemental Figure 4. SCC VII neoantigen pool identification.** C3H/HeJ mice immunized with $1\times10^7$ irradiated SCC VII cells, 50 µg polyI:C, or both in combination and subsequently challenged with $5\times10^5$ live SCC VII-Luc/GFP cells 14 days later. (**A** and **B**) Mice assessed for the presence of IFN-γ-producing splenic and Ig LN mononuclear cells at day 28 via ELISPOT after re-stimulation with NeoAg peptide pool-pulsed BMDCs. Positive controls include tripartite culturing with $2\times10^4$ irradiated/IFN-γ-treated and washed SCC VII cells or addition of ConA (n = 3 per group). All experiments were performed two or more times and data indicate means ± s.e.m.; (**A**) $*P < 0.05$, $**P < 0.01$, and $****P < 0.0001$ (Student's t test of data with SI > 2 and Poisson < 5%).

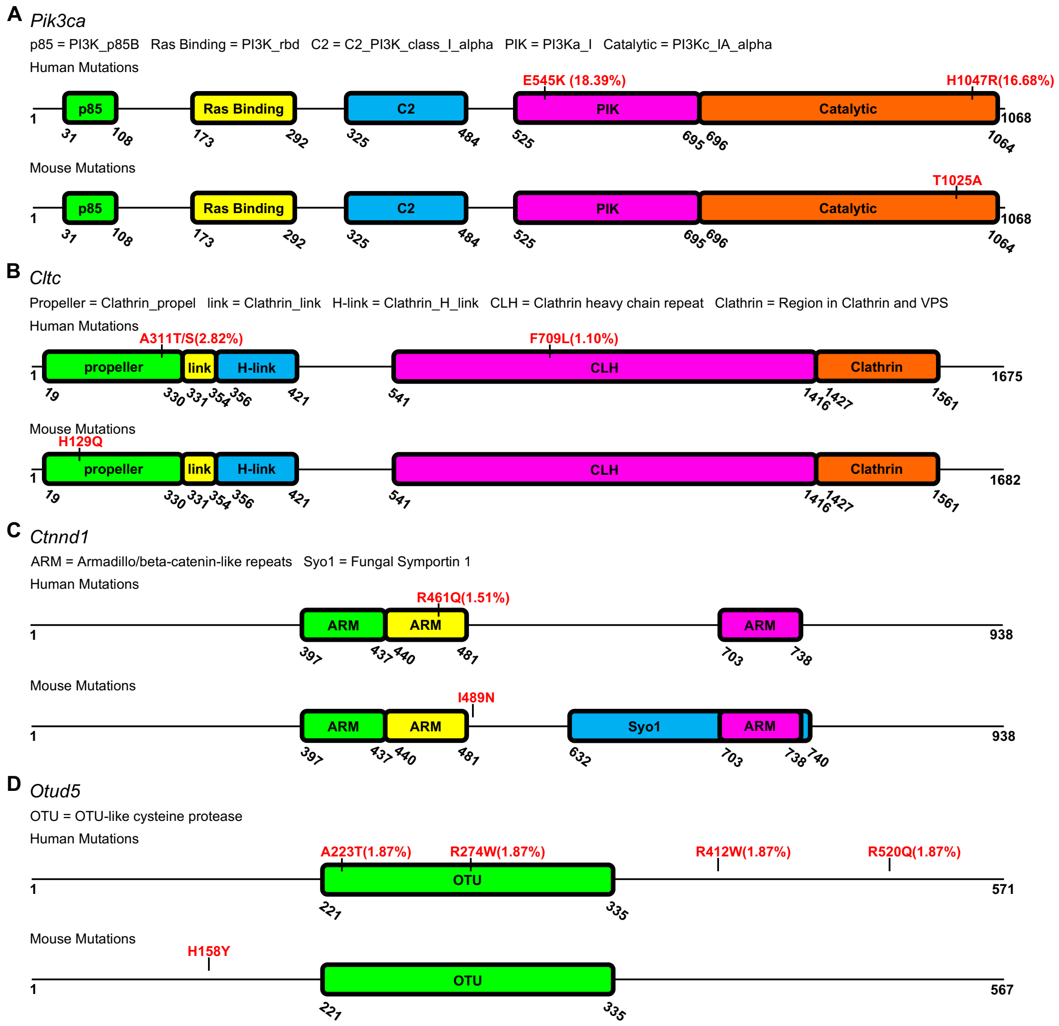


**Supplemental Figure 5. SCC VII neoantigen functional genomic positioning.** Predominant human NeoAg mutation nucleotide position and frequency in cancer patient datasets retrieved from The Cancer Genome Atlas (TCGA) database compared to C3H/HeJ mouse mutations in this study that elicited NeoAg-reactive IFN-γ ELISPOT T cell responses including (**A**) *Pik3ca*, (**B**) *Cltc*, (**C**) *Ctnnd1*, and (**D**) *Otud5*. Major functional domains are cited and aligned between murine and human genes.

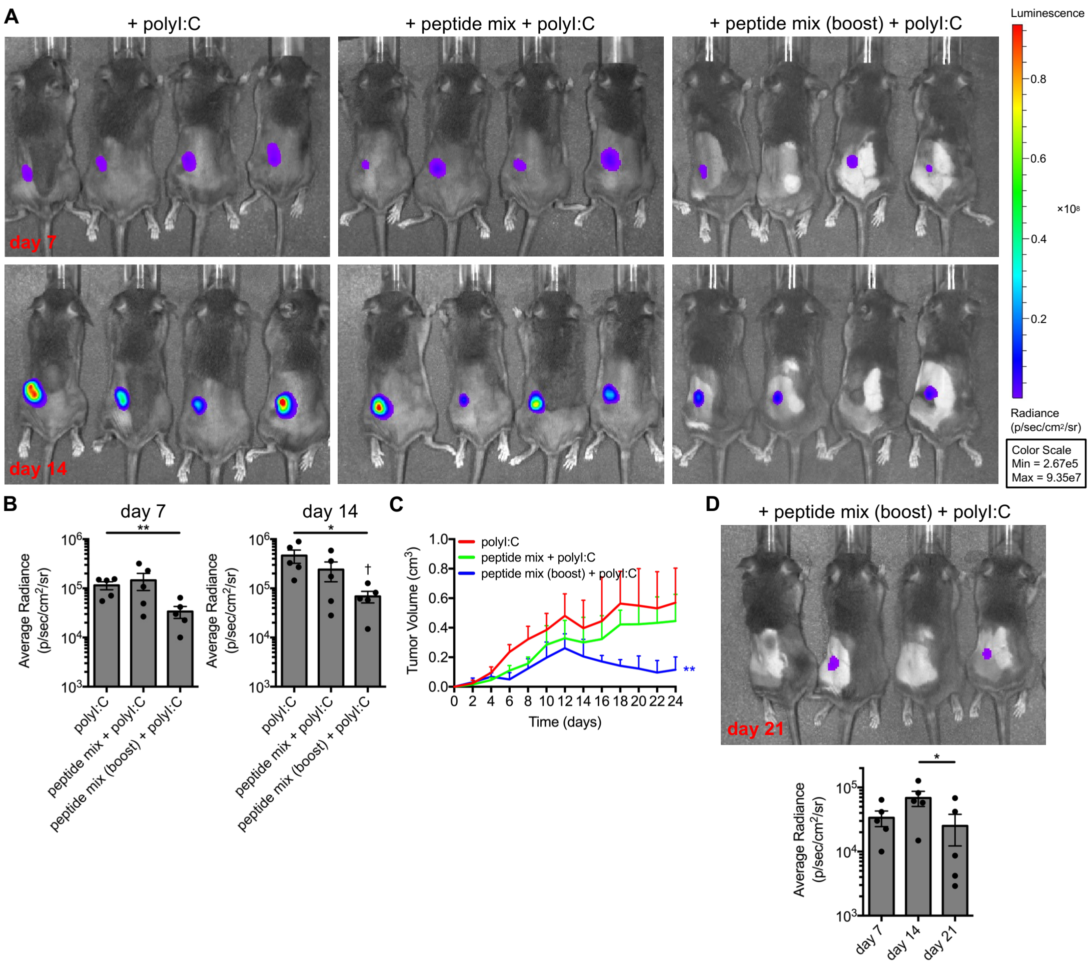


**Supplemental Figure 6. Neoantigen peptide vaccination induction of strong anti-tumor responses.** C3H/HeJ mice vaccinated with 50 μg polyI:C alone or in combination with a 5×5 μg mixture of solubilized Mut_44, Mut_48, Mut_61, Mut_65, and Mut_67 long peptides at one time point or as a booster 21 days later. All sets of mice were challenged with $5 \times 10^5$ live SCC VII-Luc/GFP cells 31 days after primary vaccination. (**A** and **B**) Bioluminescence of mice bearing SCC VII-Luc/GFP tumors at 7 and 14 days after challenge and (**C**) tumor volume kinetics (n = 6 per group). (**D**) Bioluminescence of tumors tracked to 21 days in surviving mice of the booster cohort (n = 5 per group). All experiments were performed two or more times and data indicate means ± s.e.m.; (**B** and **D**) $^{*}P < 0.05$ and $^{**}P < 0.01$ (Student's t test); (**B**) $^{\dagger}P < 0.05$ (one-way

ANOVA and Dunnett's post hoc test relative to polyI:C); (**C**) $^{**}P < 0.01$ (two-way ANOVA and Dunnett's post hoc test relative to polyI:C).

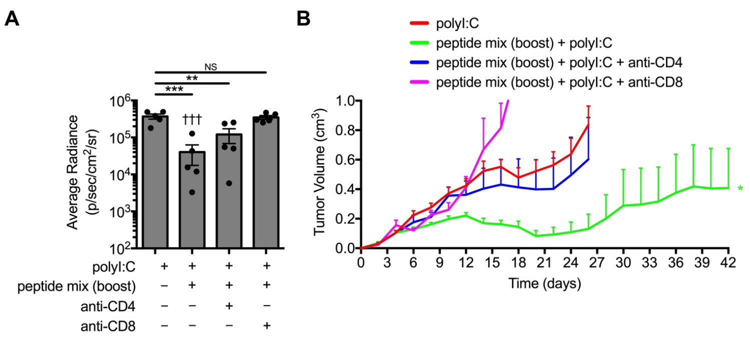


**Supplemental Figure 7. CD4$^{+}$ and CD8$^{+}$ T cells are critical during neoantigen peptide vaccination to clear SCC VII tumors in vivo.** C3H/HeJ mice vaccinated with 50 µg polyI:C alone or in combination with prime/boost regimens of a 5×5 µg mixture containing solubilized Mut_44, Mut_48, Mut_61, Mut_65, and Mut_67 long peptides. All groups of mice were challenged with $5\times10^{5}$ live SCC VII-Luc/GFP cells 31 days after primary vaccination. Depletion of CD4$^{+}$ and CD8$^{+}$ cells was conducted 5 days prior to vaccination and ceased 24 days before live tumor challenge. (**A**) Bioluminescence of mice bearing SCC VII-Luc/GFP tumors at 14 days and (**B**) tumor volume kinetics tracked to day 42 post-challenge (n = 6 per group). All experiments were performed two or more times and data indicate means ± s.e.m.; (**A**) $^{**}P < 0.01$ and $^{***}P < 0.001$ (Student's t test); $^{†††}P < 0.001$ (one-way ANOVA and Dunnett's post hoc test relative to polyI:C); (**B**) $^{*}P < 0.05$ (two-way ANOVA and Dunnett's post hoc test relative to polyI:C).

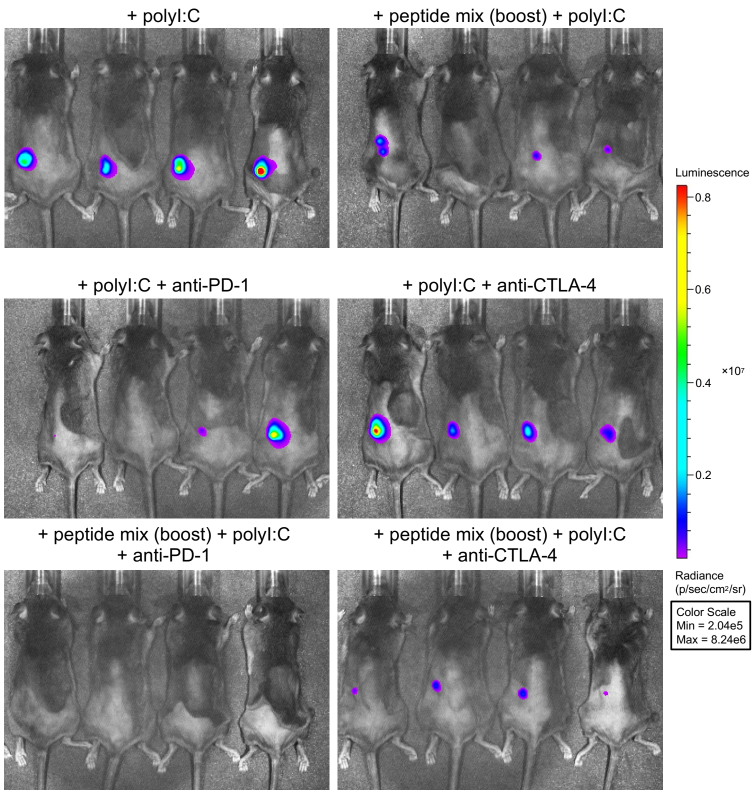


**Supplemental Figure 8. Synergy between neoantigen peptide mix and blockade of PD-1/PD-L1.** C3H/HeJ mice vaccinated with 50 µg polyI:C alone or in combination with prime/boost regimens of a 5×5 µg mixture containing solubilized Mut_44, Mut_48, Mut_61, Mut_65, and Mut_67 long peptides. All groups of mice were challenged with $5\times10^5$ live SCC VII-Luc/GFP cells 31 days after primary vaccination. For checkpoint blockade, mice received anti-PD-1 or anti-CTLA-4 therapeutically beginning at day 3 after tumor cell inoculation. Representative bioluminescence images of mice bearing live tumors at 14 days after challenge (n = 6 per group). All experiments were performed two or more times.

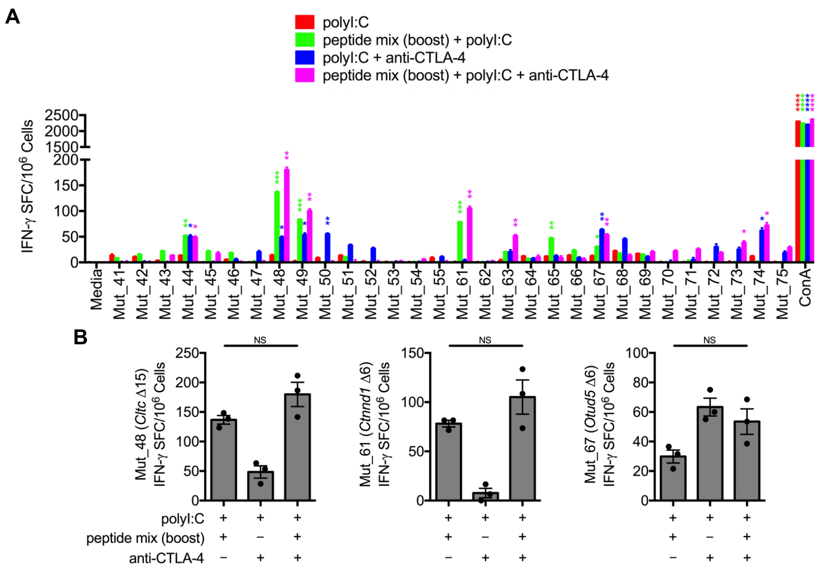


**Supplemental Figure 9. Therapeutic delivery of anti-CTLA-4 does not significantly increase *Cltc* Δ15-specific memory frequency and induces weak intermolecular epitope spreading.** C3H/HeJ mice vaccinated with 50 µg polyI:C alone or in combination with prime/boost regimens of a 5×5 µg mixture containing solubilized Mut_44, Mut_48, Mut_61, Mut_65, and Mut_67 long peptides. All groups of mice were challenged with $5 \times 10^5$ live SCC VII-Luc/GFP cells 31 days after primary vaccination. For checkpoint blockade, mice received anti-CTLA-4 therapeutically beginning at day 3 after tumor cell inoculation as 5 consecutive injections spaced 3 days apart until day 15. (**A** and **B**) Splenic and Ig LN mononuclear cells isolated at day 42 from anti-CTLA-4-treated groups and controls assessed for IFN-γ-production via ELISPOT after re-stimulation with NeoAg-pulsed BMDCs (n = 3 per group). All experiments were performed two or more times and data indicate means ± s.e.m.; (**A**) **P* < 0.05, ***P* < 0.01, ****P* < 0.001, and *****P* < 0.0001 (Student's t test of data with SI > 2 and Poisson < 5%); (**B**) NS (Student's t test).

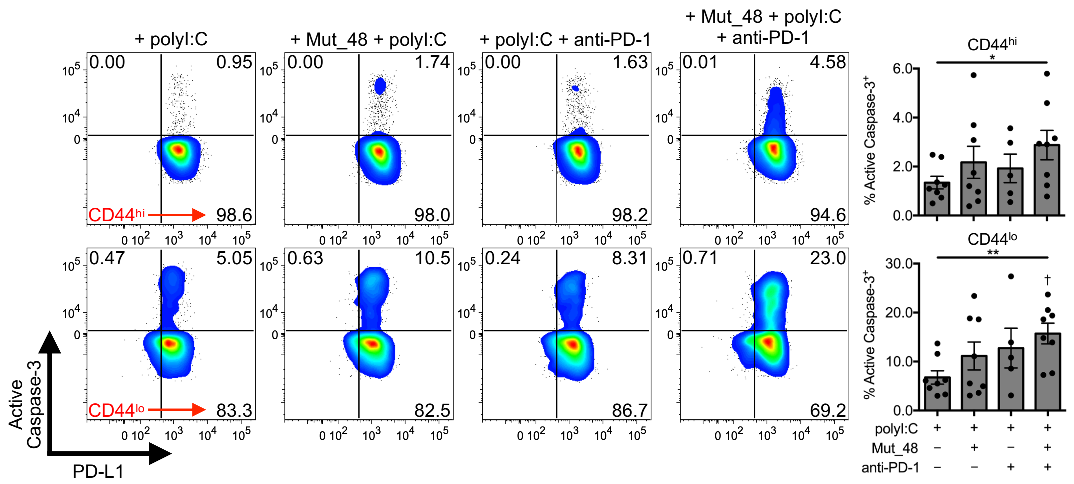


**Supplemental Figure 10. Neoantigen targeting of CD44$^{lo}$ versus CD44$^{hi}$ SCC VII populations for apoptosis.** C3H/HeJ mice injected with $5 \times 10^5$ live SCC VII-Luc/GFP cells and given 50 µg polyI:C alone or in combination with 5 µg Mut_48 peptide at day 10 post-challenge. Select groups of mice also received anti-PD-1 at days 10, 13, and 16. Frequency of active caspase-3 7 days after single day 10 immunizations (n = 5-8 per group). All experiments were performed two or more times and data indicate means ± s.e.m.; *$P < 0.05$ and **$P < 0.01$ (Student's t test); $^{\dagger}P < 0.05$ (one-way ANOVA and Dunnett's post hoc test relative to polyI:C and saline, respectively).

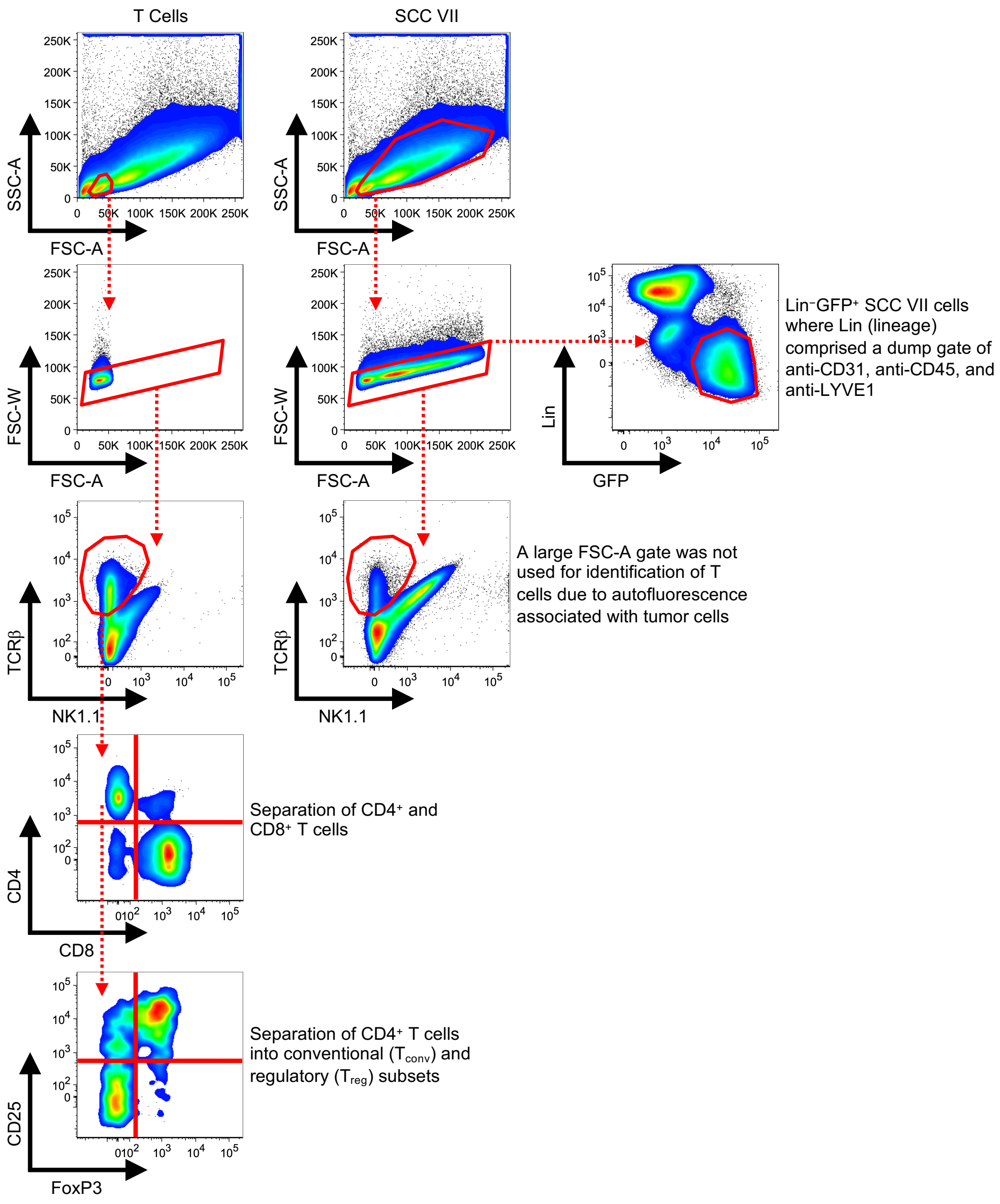


**Supplemental Figure 11. Flow cytometry gating strategy.** The complete gating strategy used in this study for identification of CD4$^{+}$ T$_{conv}$, CD4$^{+}$ T$_{reg}$, and CD8$^{+}$ CTL as well as the Lin$^{-}$GFP$^{+}$ SCC VII cell subset is graphically displayed.

| Mut_# | Peptide Pool_# | Gene Name | 20-mer Reference Peptide | 20-mer Mutated Peptide | Mutated Position in Peptide | Ref Seq | Alt Seq | VAF |
|---|---|---|---|---|---|---|---|---|
| 1 | 1 | *Hnrnpk* | SAIDTWSPSEWQMA**Y**EPQGG | SAIDTWSPSEWQMA**H**EPQGG | 15 | T | C | 0.44 |
| 2 | 1 | *Hnrnpk* | EWQMA**Y**EPQGGSGYDYSYAG | EWQMA**H**EPQGGSGYDYSYAG | 6 | T | C | 0.44 |
| 3 | 1 | *Ank2* | EGTKPCLQTPVTSE**R**GSPIV | EGTKPCLQTPVTSE**C**GSPIV | 15 | C | T | 0.24 |
| 4 | 1 | *Ank2* | PVTSE**R**GSPIVQEPEEASEP | PVTSE**C**GSPIVQEPEEASEP | 6 | C | T | 0.24 |
| 5 | 1 | *Pkp1* | TSGRQRVQEQVMMT**V**KRQKS | TSGRQRVQEQVMMT**G**KRQKS | 15 | T | G | 0.9 |
| 6 | 2 | *Pkp1* | QVMMT**V**KRQKSKSSQSSTLS | QVMMT**G**KRQKSKSSQSSTLS | 6 | T | G | 0.9 |
| 7 | 2 | *Utrn* | LQQTNSEKILLSWV**R**QTTRP | LQQTNSEKILLSWV**W**QTTRP | 15 | C | T | 0.67 |
| 8 | 2 | *Utrn* | LLSWV**R**QTTRPYSQVNVLNF | LLSWV**W**QTTRPYSQVNVLNF | 6 | C | T | 0.67 |
| 9 | 2 | *Brd8* | QLIMQTSESGISAK**S**LRGRD | QLIMQTSESGISAK**R**LRGRD | 15 | A | C | 0.53 |
| 10 | 2 | *Brd8* | GISAK**S**LRGRDSTRKQDASE | GISAK**R**LRGRDSTRKQDASE | 6 | A | C | 0.53 |
| 11 | 3 | *Gnb2l1* | PQRALRGHSHFVSD**V**VISSD | PQRALRGHSHFVSD**A**VISSD | 15 | T | C | 0.33 |
| 12 | 3 | *Gnb2l1* | HFVSD**V**VISSDGQFALSGSW | HFVSD**A**VISSDGQFALSGSW | 6 | T | C | 0.33 |
| 13 | 3 | *Shroom4* | EICPALLKRNLLPK**C**HNCRC | EICPALLKRNLLPK**Y**HNCRC | 15 | G | A | 0.33 |
| 14 | 3 | *Shroom4* | NLLPK**C**HNCRCHHHQCIRCT | NLLPK**Y**HNCRCHHHQCIRCT | 6 | G | A | 0.33 |
| 15 | 3 | *Plekhh1* | PLWRHPMLCYSQEG**L**CASLT | PLWRHPMLCYSQEG**V**CASLT | 15 | C | G | 0.7 |
| 16 | 4 | *Plekhh1* | YSQEG**L**CASLTTLPSEALQT | YSQEG**V**CASLTTLPSEALQT | 6 | C | G | 0.7 |
| 17 | 4 | *Msn* | PWSEIRNISFNDKK**F**VIKPI | PWSEIRNISFNDKK**L**VIKPI | 15 | T | C | 0.65 |
| 18 | 4 | *Msn* | FNDKK**F**VIKPIDKKAPDFVF | FNDKK**L**VIKPIDKKAPDFVF | 6 | T | C | 0.65 |
| 19 | 4 | *Pcbp1* | VEGSSGRQVTITGS**A**ASISL | VEGSSGRQVTITGS**P**ASISL | 15 | G | C | 0.35 |
| 20 | 4 | *Pcbp1* | TITGS**A**ASISLAQYLINARL | TITGS**P**ASISLAQYLINARL | 6 | G | C | 0.35 |
| 21 | 5 | *Ptpn11* | ESIVDAGPVVVHCS**A**GIGRT | ESIVDAGPVVVHCS**T**GIGRT | 15 | G | A | 0.31 |
| 22 | 5 | *Ptpn11* | VVHCS**A**GIGRTGTFIVIDIL | VVHCS**T**GIGRTGTFIVIDIL | 6 | G | A | 0.31 |
| 23 | 5 | *Epg5* | MALSSFFVEVLMMM**N**NAAVP | MALSSFFVEVLMMM**D**NAAVP | 15 | A | G | 0.33 |
| 24 | 5 | *Epg5* | VLMMM**N**NAAVPTAEFLAVSI | VLMMM**D**NAAVPTAEFLAVSI | 6 | A | G | 0.33 |
| 25 | 5 | *Chuk* | IMELQKSPYGRRQG**D**LMESL | IMELQKSPYGRRQG**N**LMESL | 15 | G | A | 0.54 |
| 26 | 6 | *Chuk* | GRRQG**D**LMESLEQRAIDLYK | GRRQG**N**LMESLEQRAIDLYK | 6 | G | A | 0.54 |
| 27 | 6 | *Hivep1* | SLSRPNSFDKPEPL**E**GGITF | SLSRPNSFDKPEPL**D**GGITF | 15 | A | C | 0.27 |
| 28 | 6 | *Hivep1* | KPEPL**E**GGITFSLQELGRTG | KPEPL**D**GGITFSLQELGRTG | 6 | A | C | 0.27 |
| 29 | 6 | *Lonp1* | CTIVTALLSLALGQ**P**VLQNL | CTIVTALLSLALGQ**A**VLQNL | 15 | C | G | 0.4 |
| 30 | 6 | *Lonp1* | LALGQ**P**VLQNLAMTGEVSLT | LALGQ**A**VLQNLAMTGEVSLT | 6 | C | G | 0.4 |
| 31 | 7 | *Dennd2a* | QRLVNVKSRLKQAP**R**YSSLD | QRLVNVKSRLKQAP**Q**YSSLD | 15 | G | A | 0.3 |
| 32 | 7 | *Dennd2a* | LKQAP**R**YSSLDRDLIEYQER | LKQAP**Q**YSSLDRDLIEYQER | 6 | G | A | 0.3 |
| 33 | 7 | *Gtf3c1* | PFVTWQVVRDILHA**T**FEESL | PFVTWQVVRDILHA**N**FEESL | 15 | C | A | 0.47 |
| 34 | 7 | *Gtf3c1* | DILHA**T**FEESLDKTSHSVGR | DILHA**N**FEESLDKTSHSVGR | 6 | C | A | 0.47 |
| 35 | 7 | *Ndst2* | DQMRLNKQFALEHG**I**PTDLG | DQMRLNKQFALEHG**L**PTDLG | 15 | A | C | 0.21 |
| 36 | 8 | *Ndst2* | ALEHG**I**PTDLGYAVAPHHSG | ALEHG**L**PTDLGYAVAPHHSG | 6 | A | C | 0.21 |
| 37 | 8 | *Dnajc22* | SMDNFYIVGLPLAV**G**LGVLL | SMDNFYIVGLPLAV**S**LGVLL | 15 | G | A | 0.21 |
| 38 | 8 | *Dnajc22* | LPLAV**G**LGVLLVAAVGNQTS | LPLAV**S**LGVLLVAAVGNQTS | 6 | G | A | 0.21 |
| 39 | 8 | *Rai14* | SEADQQDLLVLLQA**K**VASLT | SEADQQDLLVLLQA**N**VASLT | 15 | A | C | 0.4 |
| 40 | 8 | *Rai14* | VLLQA**K**VASLTLHNKELQDK | VLLQA**N**VASLTLHNKELQDK | 6 | A | C | 0.4 |
| 41 | 9 | *Dhx32* | EIRKQLLGSSPSGK**L**FCLYT | EIRKQLLGSSPSGK**R**FCLYT | 15 | T | G | 0.38 |
| 42 | 9 | *Dhx32* | SPSGK**L**FCLYTEEFASKDMR | SPSGK**R**FCLYTEEFASKDMR | 6 | T | G | 0.38 |
| 43 | 9 | *Pik3ca* | PELQSFDDIAYIRK**T**LALDK | PELQSFDDIAYIRK**A**LALDK | 15 | A | G | 0.41 |
| 44 | 9 | *Pik3ca* | AYIRK**T**LALDKTEQEALEYF | AYIRK**A**LALDKTEQEALEYF | 6 | A | G | 0.41 |
| 45 | 9 | *Dedd2* | KRTEDSCRRRRQAS**S**SSDSP | KRTEDSCRRRRQAS**R**SSDSP | 15 | T | G | 0.2 |
| 46 | 10 | *Dedd2* | RRQAS**S**SSDSPQSQWDTGES | RRQAS**R**SSDSPQSQWDTGES | 6 | T | G | 0.2 |
| 47 | 10 | *Dedd2* | RRQAS**S**SSDSPQSQWDTGSP | RRQAS**R**SSDSPQSQWDTGSP | 6 | T | G | 0.2 |
| 48 | 10 | *Cltc* | SLNTVALVTDNAVY**H**WSMEG | SLNTVALVTDNAVY**Q**WSMEG | 15 | C | G | 0.21 |
| 49 | 10 | *Cltc* | DNAVY**H**WSMEGESQPVKMFD | DNAVY**Q**WSMEGESQPVKMFD | 6 | C | G | 0.21 |
| 50 | 10 | *Aplp2* | NYLAALQSDPPRPH**R**ILQAL | NYLAALQSDPPRPH**H**ILQAL | 15 | G | A | 0.3 |
| 51 | 11 | *Aplp2* | PPRPH**R**ILQALRRYVRAENK | PPRPH**H**ILQALRRYVRAENK | 6 | G | A | 0.3 |
| 52 | 11 | *Flna* | VDSLTKVATVPQHA**T**SGPGP | VDSLTKVATVPQHA**P**SGPGP | 15 | A | C | 0.76 |
| 53 | 11 | *Flna* | VPQHA**T**SGPGPADVSKVVAK | VPQHA**P**SGPGPADVSKVVAK | 6 | A | C | 0.76 |
| 54 | 11 | *Cep68* | YPLRPGPQLPKQPE**S**HVLTE | YPLRPGPQLPKQPE**C**HVLTE | 15 | A | T | 0.23 |
| 55 | 11 | *Cep68* | PKQPE**S**HVLTEPVLQDSGVD | PKQPE**C**HVLTEPVLQDSGVD | 6 | A | T | 0.23 |

Continued on next page

| | | | | | | | | |
|---|---|---|---|---|---|---|---|---|
| 56 | 12 | *Usp7* | PKGDGKWCKFDDDVVSRCTK | PKGDGKWCKFDDDVLSRCTK | 15 | G | C | 0.34 |
| 57 | 12 | *Usp7* | FDDDVVSRCTKEEAIEHNYG | FDDDVLSRCTKEEAIEHNYG | 6 | G | C | 0.34 |
| 58 | 12 | *Phb2* | TALKLLLGAGAVAYGVRESV | TALKLLLGAGAVAYDVRESV | 15 | G | A | 0.31 |
| 59 | 12 | *Phb2* | GAVAYGVRESVFTVEGGHRA | GAVAYDVRESVFTVEGGHRA | 6 | G | A | 0.31 |
| 60 | 12 | *Ctnnd1* | TLWNLSSHDSIKMEIVDHAL | TLWNLSSHDSIKMENVDHAL | 15 | T | A | 0.29 |
| 61 | 13 | *Ctnnd1* | SIKMEIVDHALHALTDEVII | SIKMENVDHALHALTDEVII | 6 | T | A | 0.29 |
| 62 | 13 | *Hnrnpdl* | LIDPKRAKALKGKEPPKKVF | LIDPKRAKALKGKEHPKKVF | 15 | C | A | 0.36 |
| 63 | 13 | *Hnrnpdl* | LKGKEPPKKVFVGGLSPDTS | LKGKEHPKKVFVGGLSPDTS | 6 | C | A | 0.36 |
| 64 | 13 | *Otud5* | MDPATVEQEHWFEKALRDKK | MDPATVEQEYWFEKALRDKK | 10 | C | T | 0.55 |
| 65 | 13 | *Otud5* | RIEAMDPATVEQQEHWFEKA | RIEAMDPATVEQQEYWFEKA | 15 | C | T | 0.55 |
| 66 | 14 | *Otud5* | TVEQEHWFEKALRDKKGFII | TVEQEYWFEKALRDKKGFII | 6 | C | T | 0.55 |
| 67 | 14 | *Otud5* | VEQQEHWFEKALRDKKGFII | VEQQEYWFEKALRDKKGFII | 6 | C | T | 0.55 |
| 68 | 14 | *Lamb1* | TNATKEKVDKSNEDLRNLIK | TNATKEKVDKSNEDPRNLIK | 15 | T | C | 0.45 |
| 69 | 14 | *Lamb1* | KSNEDLRNLIKQIRNFLTED | KSNEDPRNLIKQIRNFLTED | 6 | T | C | 0.45 |
| 70 | 14 | *Smg1* | LIQWVDGATPLFGLYKRWQQ | LIQWVDGATPLFGLSKRWQQ | 15 | A | C | 0.34 |
| 71 | 15 | *Smg1* | PLFGLYKRWQQREAALQAQK | PLFGLSKRWQQREAALQAQK | 6 | A | C | 0.34 |
| 72 | 15 | *Slc26a11* | TVIVGLGELLEDFQKKGVAL | TVIVGLGELLEDFQQKGVAL | 15 | A | C | 0.33 |
| 73 | 15 | *Slc26a11* | LEDFQKKGVALAFVGLQVPV | LEDFQQKGVALAFVGLQVPV | 6 | A | C | 0.33 |
| 74 | 15 | *Med13* | CTHILVFPTSASVQVASATY | CTHILVFPTSASVQGASATY | 15 | T | G | 0.35 |
| 75 | 15 | *Med13* | SASVQVASATYTTENLDLAF | SASVQGASATYTTENLDLAF | 6 | T | G | 0.35 |
| 76 | 16 | *Vps25* | NKSSFLIMWRRPEEWGKLIY | NKSSFLIMWRRPEELGKLIY | 15 | G | T | 0.34 |
| 77 | 16 | *Vps25* | RRPEEWGKLIYQWVSRSGQN | RRPEELGKLIYQWVSRSGQN | 6 | G | T | 0.34 |
| 78 | 16 | *Zfp820* | LENYNNLFFVETHGMCPKYK | LENYNNLFFVETHGICPKYK | 15 | G | A | 0.58 |
| 79 | 16 | *Zfp820* | VETHGMCPKYKNILDQDLQH | VETHGICPKYKNILDQDLQH | 6 | G | A | 0.58 |
| 80 | 16 | *Tpcn1* | YLSIELYFIMNLLLAVVFDT | YLSIELYFIMNLLLVVVFDT | 15 | C | T | 0.32 |
| 81 | 16 | *Tpcn1* | MNLLLAVVFDTFNDIEKHKF | MNLLLVVVFDTFNDIEKHKF | 6 | C | T | 0.32 |

**Supplemental Table 1. Selected neoantigen peptides.** Sequences of selected nonsynonymous missense mutations and reference 20-mers used in ELISPOT and in vivo experiments. NeoAg identified after Exome-Seq of SCC VII tumor compared to the mm10 reference genome. Amino acid mutations indicated (red). Expression of exome variants confirmed by aligning exome variant results with RNA-Seq data at a VAF > 0.20 cutoff.

| Mutation | Color |
|---|---|
| frameshift variant | dark purple |
| missense variant & splice region variant | red |
| missense variant & in-frame deletion | red |
| intron variant | grey |
| 3'-UTR variant | orange |
| in-frame deletion | light blue |
| frameshift variant & splice donor variant & splice region variant & intron variant | light purple |
| downstream gene variant | light green |
| non-coding exon variant | light grey |
| 5'-UTR variant | light orange |
| synonymous variant | yellow |
| disruptive in-frame deletion | light blue |
| in-frame insertion | dark blue |
| intergenic region | black |
| in-frame insertion & synonymous variant | light blue |
| sequence feature | very light blue |
| stop gained | dark green |
| upstream gene variant | light green |
| splice region variant & synonymous variant | yellow green |
| splice region variant & intron variant | light pink red |

**Supplemental Table 2. Circos plot legend.** Displayed are the color-coded and annotated variants displayed in the Figure 1G somatic mutation Circos plot (level 2) of the main text identified after Exome-Seq of SCC VII tumor compared to the mm10 reference genome.

| Antibody | Clone | Reactivity | Fluorochrome, Dilution | Manufacturer |
|---|---|---|---|---|
| anti-Active Caspase-3 | 5A1E | Mouse/Human | PE, 1:50 | Cell Signaling Technology |
| anti-ALDH1A1 | 03 | Human | PE, 1:50 | Sino Biological |
| anti-ALDH1A1 | EP1933Y | Mouse | AF488, 1:200 | Abcam |
| anti-CD4 | RM4-5 | Mouse | APC-eF780, 1:400 | eBioscience |
| anti-CD4 | RM4-5 | Mouse | BUV737, 1:400 | BD Biosciences |
| anti-CD4 | RM4-5 | Mouse | PerCP-Cy5.5, 1:400 | Tonbo Biosciences |
| anti-CD8α | 53-6.7 | Mouse | BV570, 1:400 | BioLegend |
| anti-CD8α | 53-6.7 | Mouse | eF450, 1:400 | eBioscience |
| anti-CD25 | 3C7 | Mouse | PE-Cy7, 1:400 | BioLegend |
| anti-CD25 | PC61.5 | Mouse | PerCP-Cy5.5, 1:400 | Tonbo Biosciences |
| anti-CD31 | 390 | Mouse | PE-Cy7, 1:400 | BioLegend |
| anti-CD44 | IM7 | Mouse/Human | AF488, 1:400 | BioLegend |
| anti-CD44 | BJ18 | Human | PerCP-Cy5.5, 1:200 | BioLegend |
| anti-CD44 | IM7 | Mouse/Human | PerCP-Cy5.5, 1:400 | eBioscience |
| anti-CD45 | 30-F11 | Mouse | BUV615, 1:400 | BD Biosciences |
| anti-CD45 | 30-F11 | Mouse | PE-Cy7, 1:400 | eBioscience |
| anti-CD62L | MEL-14 | Mouse | BB515, 1:400 | BD Biosciences |
| anti-CD69 | H1.2F3 | Mouse | SB645, 1:400 | Invitrogen |
| anti-CD127 | A7R34 | Mouse | PerCP-Cy5.5, 1:400 | BioLegend |
| anti-CX3CR1 | SA011F11 | Mouse | BV785, 1:200 | BioLegend |
| anti-EGFR | AY13 | Human | PE-Cy7, 1:200 | BioLegend |
| anti-EGFR | H11 | Mouse | AF647, 1:400 | Invitrogen |
| anti-Eomes | Dan11mag | Mouse | PE-eF610, 1:200 | Thermo Scientific |
| anti-EpCAM | 9C4 | Human | APC-Fire750, 1:200 | BioLegend |
| anti-EpCAM | G8.8 | Mouse | APC-eF780, 1:400 | eBioscience |
| anti-FoxP3 | FJK-16s | Mouse | AF532, 1:200 | Thermo Scientific |
| anti-FoxP3 | FJK-16s | Mouse | PE, 1:200 | eBioscience |
| anti-GFP | pAb | WT/rGFP/eGFP | FITC, 1:200 | Rockland Immunochemicals |
| anti-GzmB | GB11 | Mouse/Human | Pacific Blue, 1:200 | BioLegend |
| anti-ICOS | C398.4A | Mouse | AF700, 1:400 | BioLegend |
| anti-Ki-67 | B56 | Mouse/Human | BUV395, 1:200 | BD Biosciences |
| anti-KLRG1 | 2F1 | Mouse | BV605, 1:400 | BioLegend |
| anti-Lag-3 | C9B7W | Mouse | BUV496, 1:400 | BD Biosciences |
| anti-LYVE1 | ALY7 | Mouse | PE-Cy7, 1:400 | eBioscience |
| anti-NK1.1 | PK136 | Mouse | PE-Cy7, 1:400 | eBioscience |
| anti-NKp46 | 29A1.4 | Mouse | BUV805, 1:400 | BD Biosciences |
| anti-PD-1 | 29F.1A12 | Mouse | APC-Fire750, 1:400 | BioLegend |
| anti-PD-L1 | MIH5 | Mouse | APC, 1:400; PE, 1:400 | eBioscience |
| anti-SLAMF6 | 13G3 | Mouse | BV480, 1:400 | BD Biosciences |
| anti-T-bet | 4B10 | Mouse/Human | BV421, 1:200 | BioLegend |
| anti-TCF-1 | C63D9 | Mouse/Human | PE, 1:100 | Cell Signaling Technology |
| anti-TCRβ | H57-597 | Mouse | APC, 1:400; FITC, 1:400 | Tonbo Biosciences |
| anti-TCRβ | H57-597 | Mouse | BV510, 1:400 | BD Biosciences |
| anti-TCRγδ | GL3 | Mouse | PerCP-eF710, 1:400 | Thermo Scientific |
| anti-Thy1.2 | 53-2.1 | Mouse | BV750, 1:400 | BD Biosciences |
| anti-Tim-3 | RMT3-23 | Mouse | BV711, 1:400 | BioLegend |
| anti-TOX | TXRX10 | Mouse/Human | eF660, 1:200 | Thermo Scientific |

**Supplemental Table 3. Antibody list.** These antibodies were used in FACS experiments.
